# Molecular diagnosis and prognosis of cancers of unknown-primary (CUPs): progress from a microRNA-based droplet digital PCR assay

**DOI:** 10.1101/2021.02.03.429509

**Authors:** Noemi Laprovitera, Mattia Riefolo, Elisa Porcellini, Giorgio Durante, Ingrid Garajova, Francesco Vasuri, Ariane Aigelsreiter, Nadia Dandachi, Giuseppe Benvenuto, Federico Agostinis, Silvia Sabbioni, Ioana Berindan-Neagoe, Chiara Romualdi, Andrea Ardizzoni, Davide Trerè, Martin Pichler, Antonia D’Errico, Manuela Ferracin

## Abstract

Metastasis is responsible for the majority of cancer-related deaths. Particularly challenging is the management of metastatic cancer of unknown primary site (CUP), whose tissue-of-origin (TOO) remains undetermined even after expensive investigations. CUP therapy is rather unspecific and poorly effective. Molecular approaches developed to identify CUPs’ potential tissue-of-origin, can overcome some of these issues. In this study, we applied a pre-determined set of microRNAs (miRNAs) to infer the TOO of 53 metastatic cancers of unknown or uncertain origin. We designed a molecular assay to quantify 89 miRNAs at the copy number level, using EvaGreen-based Droplet Digital PCR. We assessed miRNA expression in 159 samples including primary tumors from 17 tumor classes (reference set), metastases of known and unknown origin. We applied two different statistical models for class prediction to obtain CUP’s most probable TOOs. Specifically, we used the shrunken centroids using PAMR (Prediction Analysis of Microarrays for R) and the least absolute shrinkage and selection operator (LASSO) models. The molecular test was successfully applied to FFPE samples and provided a site-of-origin identification within one-week from the biopsy procedure. The most frequently predicted origins were gastrointestinal, pancreas, breast and lung. The assay was applied to multiple metastases from the same CUP, collected from different metastatic sites: the molecular prediction revealed an impressive agreement in site-of-origin prediction, intrinsically validating our assay. The final prediction was compared with the clinico-pathological hypothesis of primary site. Moreover, a panel of 14 miRNAs proved to have prognostic value and being associated with overall survival. Our study demonstrated that miRNA expression profiling in CUP samples could be employed as diagnostic and prognostic test. Our molecular analysis can be performed on-request, concomitantly with standard diagnostic workup and in association with genetic profiling, to offer valuable indication about the possible primary site, thereby supporting treatment decisions.

## Introduction

Cancer of Unknown Primary origin (CUP) describes newly diagnosed tumors presenting as metastatic cancers, whose primary site cannot be recognized after detailed standardized physical examinations, blood analyses, imaging and immunohistochemical (IHC) testing [1]. CUP biology represents a real riddle and several theories have been proposed to describe CUP origin. According to the two prevailing hypotheses, CUPs could either be metastatic tumors that originate from an undetectable, small, dormant or later regressed primary tumors or represent metastatic entities with no existing primary that derive from cancer stem cells and early acquire the metastatic ability [1].

Post-mortem investigations on CUP patients reported the identification of a primary tumor in about 75% of cases and highlighted the prevalent epithelial origin of CUPs. The most common primary sites were represented by lung, pancreas, hepatobiliary tract, kidney, colon, genital organs and stomach [2]. Population-based studies reported decreasing trends of CUP incidence in different countries in the last decade, possibly as a consequence of novel diagnostic techniques that improved primary site identification or a more consequent and widespread approach to follow standardized diagnostic work-up guidelines [3]. Nonetheless, incidence-rates still vary among different countries worldwide.

International guidelines for tumor treatment are essentially based on primary site indication. Therefore, CUP treatment requires a rather unspecific blind approach, which is very challenging for the treating physicians. As a consequence, CUPs are usually treated with empiric platinum-based chemotherapy regimens that are poorly effective. CUP patients have a short life-expectancy (average overall survival 4-9 months, 20% survive more than 1 year) that did not improve in the last decades. In the most recent CUP NCCN guidelines (v.2/2020), there are eleven different chemotherapy regimens indicated for adenocarcinoma and nine for squamous histology. However, these regimens remain empirical since they are mostly based on single-arm phase II clinical trials [4–6] and small randomized prospective trials [7–9]. In addition, the lack of primary tumor definition, prevent most patients to be treated in clinical practice with novel very effective treatment such as immunotherapy or molecular targeted therapies for which current registered indications are mostly disease-oriented. Finally, patients with occult primary tumors suffer a great psychological burden of an unidentified disease. The use of molecular tests that could identify the most probable site-of-origin or an approach based on personalized medicine, may be useful to assist in the selection of the best treatment options and potentially improve CUPs prognosis and survival.

The identification of druggable alterations in CUP tumors could improve the otherwise limited treatment options. Recently, several studies focused on the analysis of CUP mutational profiles [10–12]. A comprehensive retrospective analysis, using the 236-gene FoundationOne assay (Roche), explored the genomic profiles of 200 CUPs [11]. At least 1 clinically relevant genetic alteration was found in 96% of CUPs, with a mean of 4.2 alterations per tumor. Most frequently mutated genes were *TP53* (55%), *KRAS* (20%), *CDKN2A* (19%), *MYC* (12%), *ARID1A* (11%) and *MCL1* (10%). According to this study, potentially druggable mutations were discovered in 20% of CUPs. Varghese et al. identified the actionable mutations in a dataset of 150 CUPs analyzed with the MSK-IMPACT panel [12] and in another dataset of 200 CUPs from Ross et al. [11]. Potentially druggable alterations were present in 30% of CUP cases (FDA level 2-3 of evidence for actionability) [12]. Few patients in Varghese study were treated with targeted therapies, overall showing benefit and achieved longer survival.

Another way to improve the choice of CUP therapeutic options, is the prediction of CUP site-of-origin using molecular assays. This strategy is based on the observation that metastatic tumor cells retain some molecular characteristics of the tissue of origin, despite their going through de-differentiation and epithelial-mesenchymal transition programs. This tissue-specific molecular signature can be leveraged to obtain an inference about CUP origin. In the past decade, several molecular classifiers were developed. These classifiers were built based on gene expression profiles (GEP) [13–16], microRNAs [17, 18] or DNA methylation [19].

A number of studies reported evidences in favor of this hypothesis, showing a prolonged survival in patients treated with cancer-specific agents compared to standard chemotherapy [20, 21, 19, 22, 23]. Results from a prospective study on nearly 300 patients with unknown primary tumors who were treated according to GEP molecular prediction revealed a significant increase in median survival time (12.5 months) [24].

In addition, GEP proved a higher diagnostic accuracy compared to standard immunohistochemistry (IHC) staining in the identification of CUP primary site, especially in moderately or poorly differentiated cases [24, 25]. The most recent NCCN CUP Guidelines [26] support the use of gene expression profiling to get a diagnostic benefit in CUP management, though the achievement of a clinical benefit still needs to be determined. Results from the phase III clinical trial NCT03278600 could help to clarify the value of tissue-of-origin profiling in predicting primary site and directing therapy in CUP patients.

However, the analysis of GEP in archival formalin-fixed, paraffin-embedded (FFPE) tissues is limited by the quality of extracted RNA, which is usually low. Thus, the reported rate of technical success of GEP assays (i.e. CancerTypeID assay) is 85% [21]. On the contrary, microRNAs (miRNAs) are robustly detected irrespective of the quality of the tissue sample [27, 28] and are highly stable and resistant to RNAase degradation either in compromised archived clinical specimens [29, 30] or biological fluids [31]. Molecular miRNA profiling of FFPE samples could be successfully obtained from all the available samples [17, 32].

Independently from the molecular assay choice, assessing the true clinical benefit of molecular profiling is challenging because it relies on surrogate measures (correlation with IHC findings, clinical presentation or response to therapy), given that a real primary site identification is seldom available. In a previous microarray-based study we identified a cancer-type specific miRNA signature able to predict metastatic tumor tissue-of-origin of CUPs among ten possible primary sites [17]. This predictive tool was employed in few occasions to provide clinicians with indications of a possible primary site [33]. However, microarray technology limitations prevent the execution of such analysis on a routine basis. To extend the analysis to more tumor types and overcome the technical limits of microarray technology, we developed a miRNA-based molecular assay for a rapid, on-demand molecular tumor characterization and primary site prediction [34]. Unlike previous assays, our test employs droplet digital PCR (ddPCR) technology to assess the absolute level of a pre-determined set of 89 miRNAs in FFPE tumor tissues. This assay is applied here to predict the most probable primary tissue(s) of a set of 53 cancers of unknown or uncertain origin, obtaining a broad spectrum of primary site predictions with different levels of confidence.

## Methods

### Patients and tumor samples

A total number of 159 formalin-fixed paraffin-embedded (FFPE) samples from 148 patients was collected for this study. Patients were diagnosed and treated at Sant’Orsola-Malpighi Bologna University Hospital, Italy (N=84), at the University Hospital of Ferrara, Italy (N=50) or at the Medical University of Graz, Austria (N=14). The study cohort consists of patients with tumors with a clearly recognized primary site (N=106) and patients with cancer of unknown or uncertain origin (CUPs, N=53). A summary of samples and patients enrolled in the study is reported in **Table 1**. Primary tumors included samples obtained from the following tumor sites/types: lung (LUAD, adenocarcinoma, N=5 and LUSC, squamous cell carcinoma, N=3), pancreas (PAAD, exocrine adenocarcinoma, N=5), liver (LIHC, hepatocellular carcinoma, N=6), biliary tract (CHOL, cholangiocarcinoma, N=6), kidney (KICA, which includes kidney renal clear cell carcinoma or KIRC, N=5 and kidney renal papillary cell carcinoma or KIRP, N=3), colorectum (CRC, adenocarcinoma, N=7), testis (TGSC, germ cell seminomatous carcinoma, N=4), endometrium (UCEC, adenocarcinoma, N=5), stomach (STAD, adenocarcinoma, N=5), bladder (BLCA, transitional cell carcinoma, N=4), breast (BRCA, non-special type and lobular carcinoma, N=5), triple negative breast cancer (TNBC, N=3), prostate (PRAD, adenocarcinoma, N=5), melanoma (SKCM, melanoma of skin, N=7), head and neck (HNSC, squamous cell carcinoma, N=6) and gastrointestinal neuroendocrine carcinoma (GI-NET, N=5). We assessed 10 metastases of known origin, derived from lung, melanoma, stomach, prostate, head and neck, kidney, colon, breast, pancreas and endometrium. A total number of 53 CUPs were included in this study, specifically 43 retrospective and 10 prospective cases. Moreover, from 5 retrospective CUP patients we were able to obtain metastatic biopsies collected from multiple sites that were independently analyzed. CUP diagnosis was obtained after detailed clinical and pathological investigations. For each sample a full IHC panel was assessed at the time of diagnosis and the outcome was recorded. However, we need to underline that our collection of CUP samples is heterogeneous since it derives from patients that received the diagnosis in different time; specifically, 26% of them received the diagnosis of CUP between 2005-2009, 49% between 2010-2014 and 24% between 2015-2019.

**Table 1.**
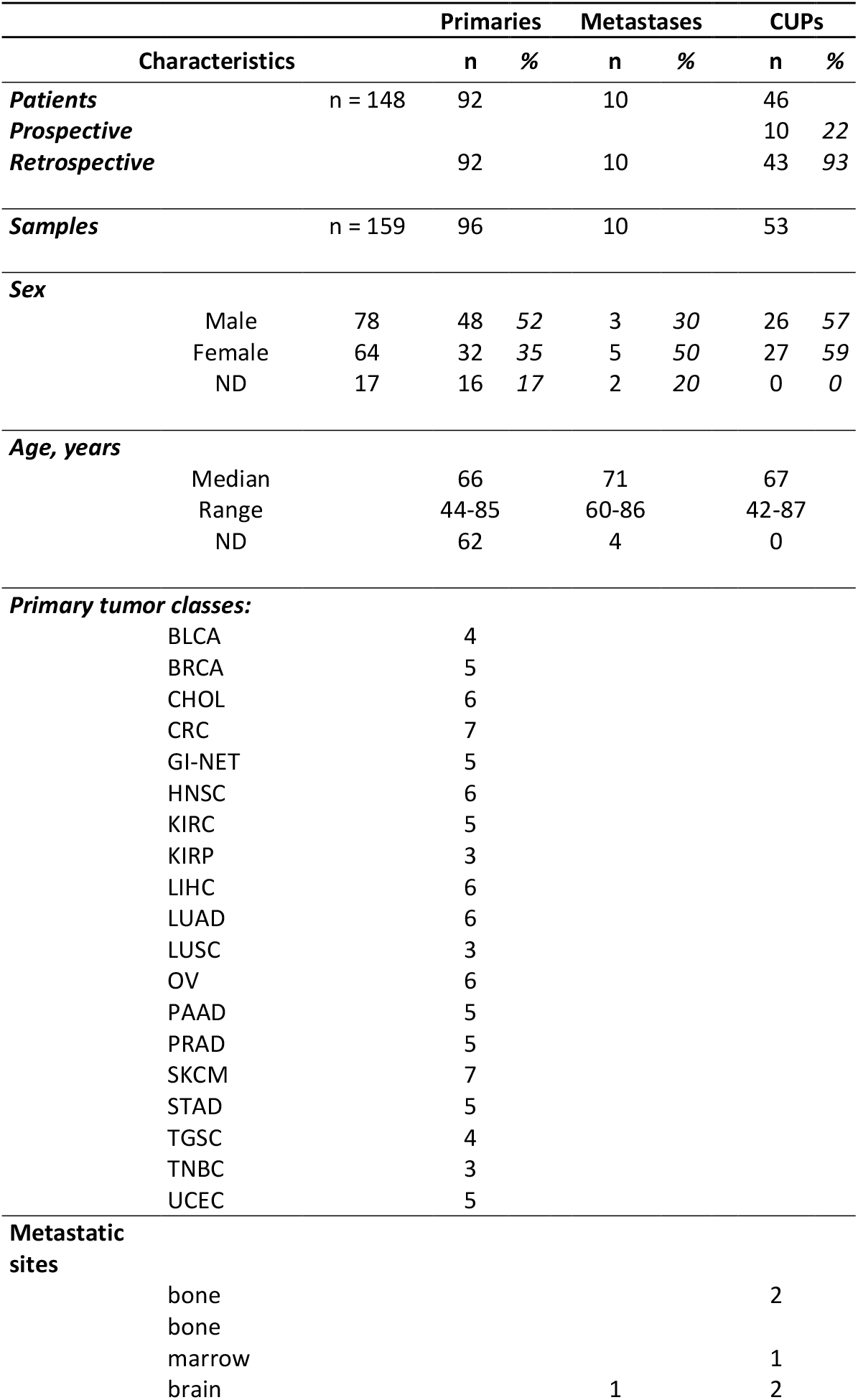
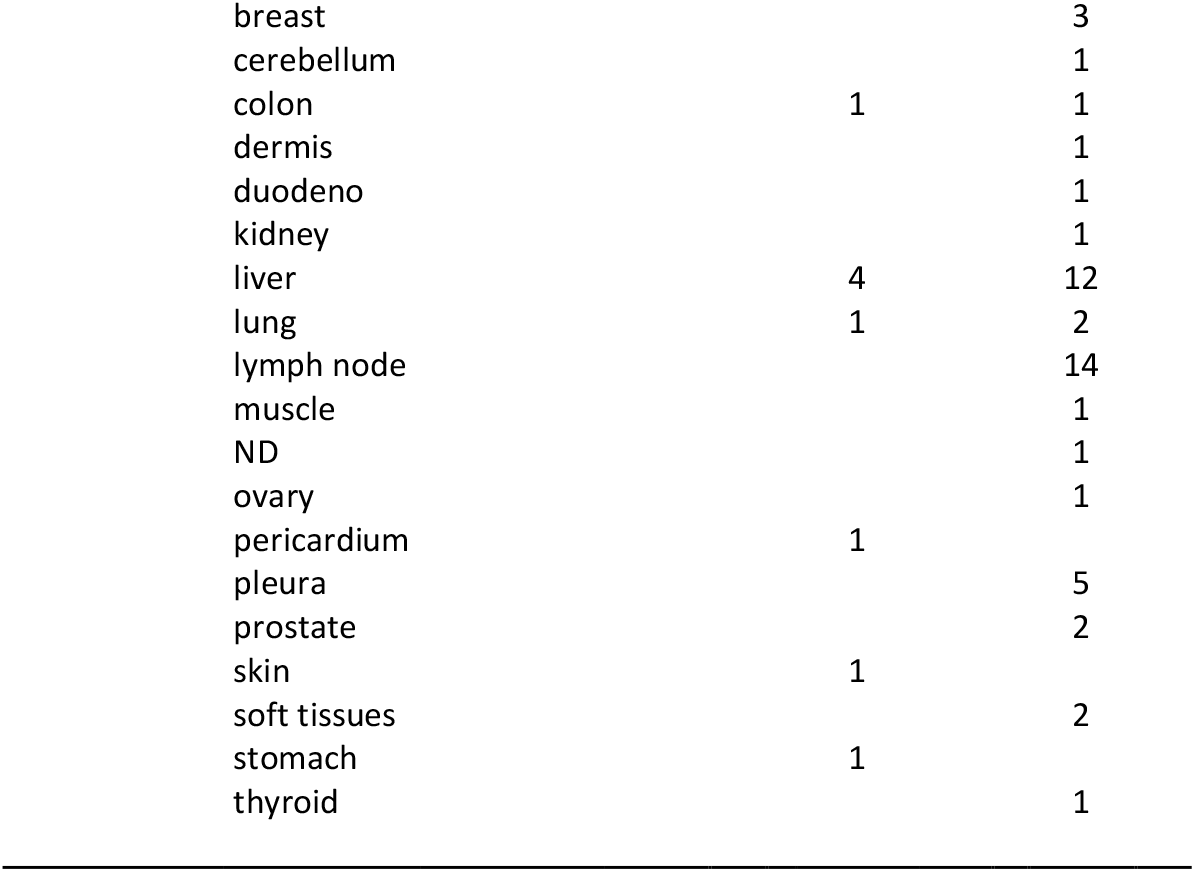
Summary of samples and patients enrolled in the study

For each sample, 10 μm thick tissue sections (N=2-5) were obtained. The first section was stained with hematoxylin-eosin (HE) and examined by an expert pathologist to select the tumor area, which was grossly dissected before RNA extraction. Tumor cell fraction was evaluated to select samples with at least 30% cellularity. The study was conducted in accordance with the Declaration of Helsinki, and the protocol was approved by the Ethics Committee Center Emilia-Romagna Region – Italy (protocol 130/2016/U/Tess) and Medical University of Graz (vote no. 30-520 ex 17/18). Prospective patients provided written informed consent. Detailed pathological characteristics of cancer patients are available in **Supplementary Table 1** and **Table 2**.

**Table 2.**
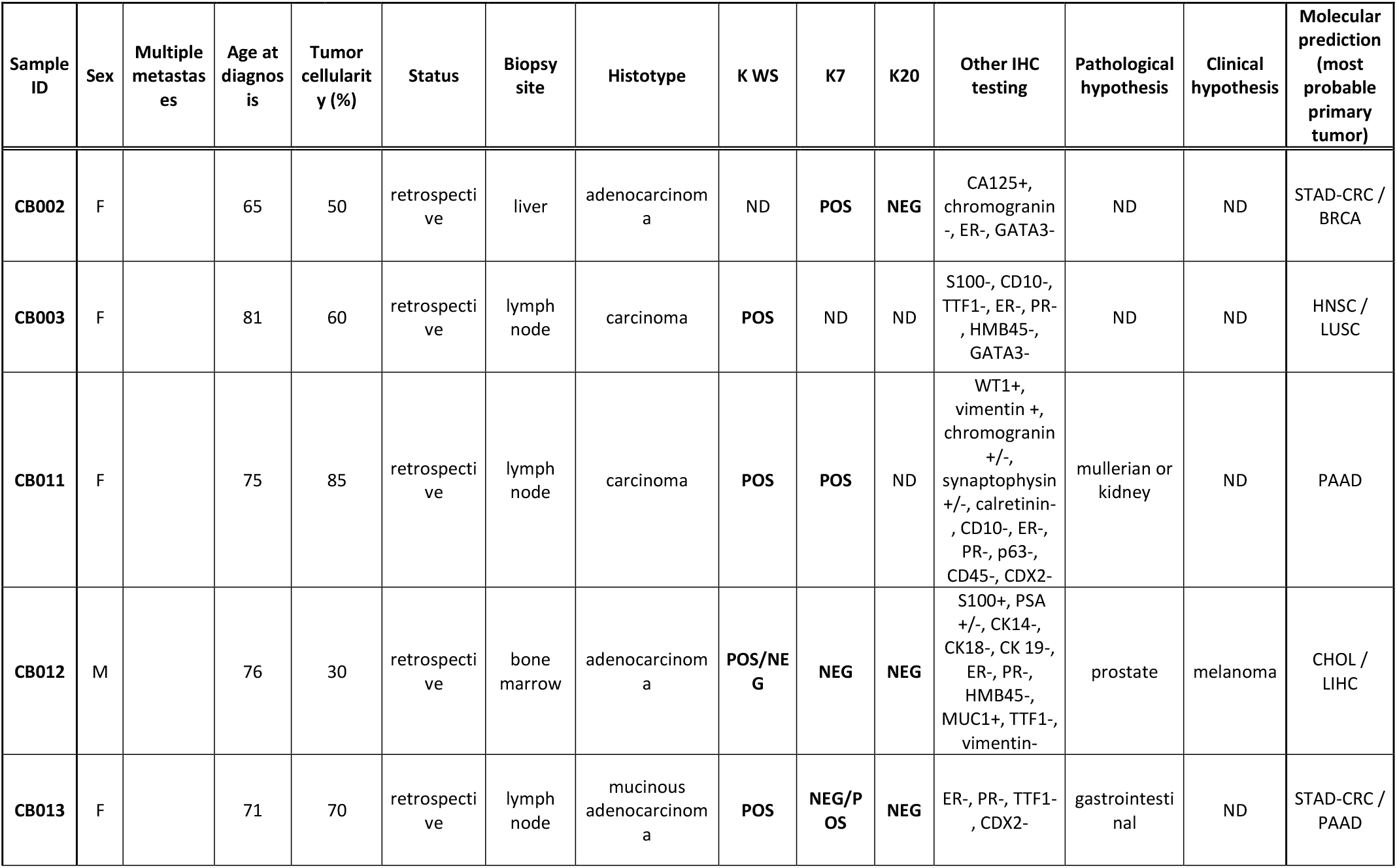
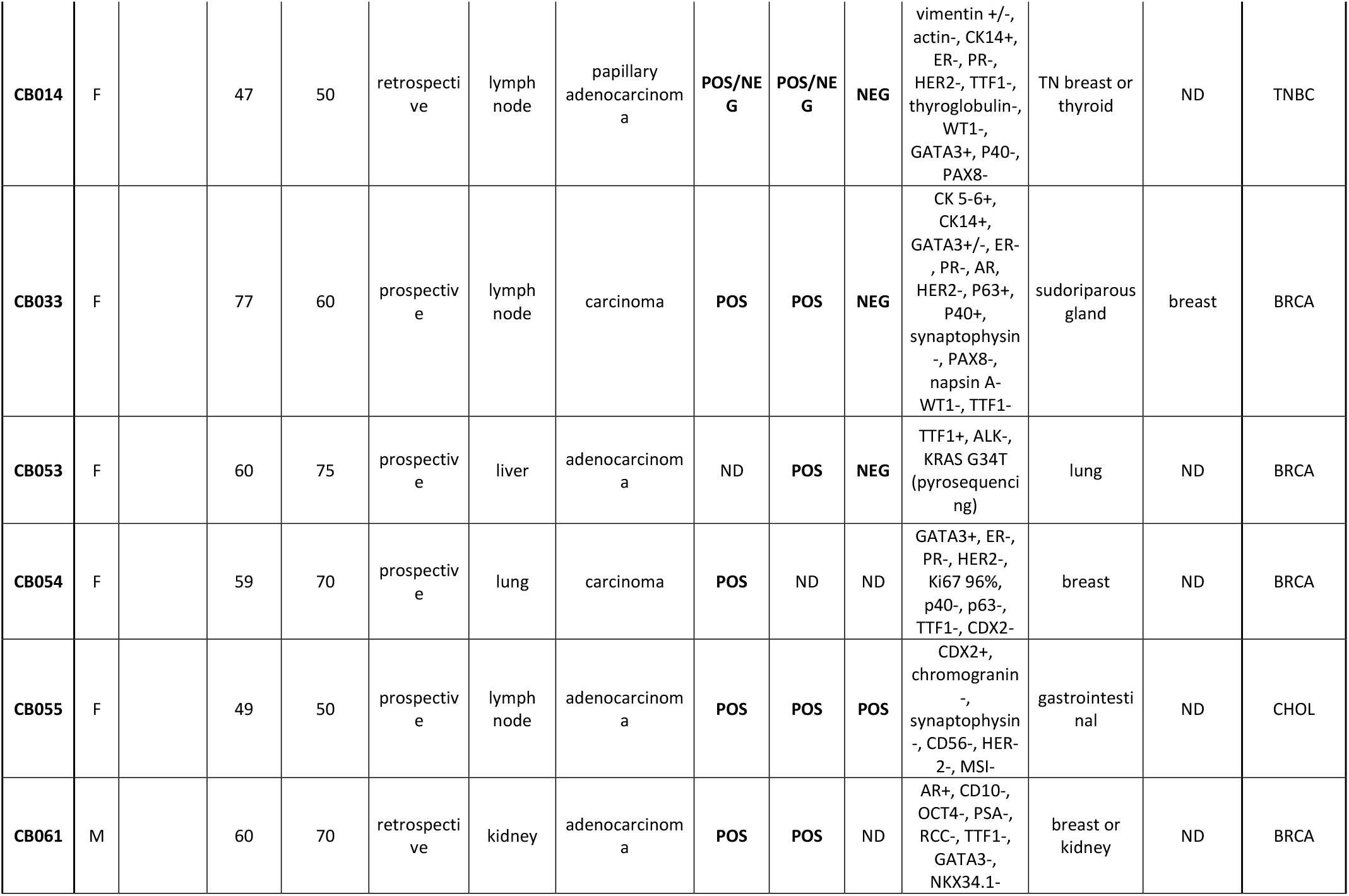
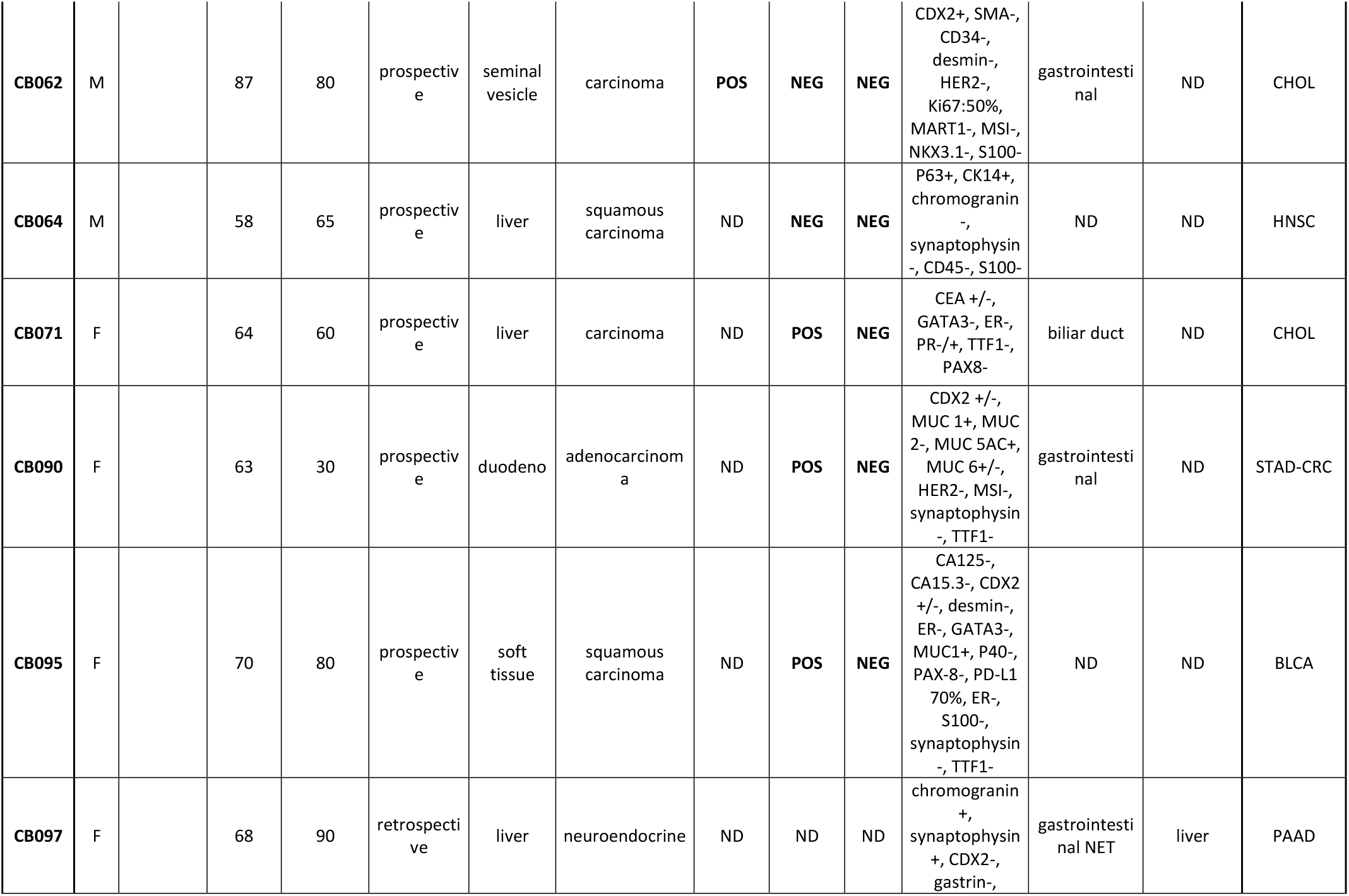
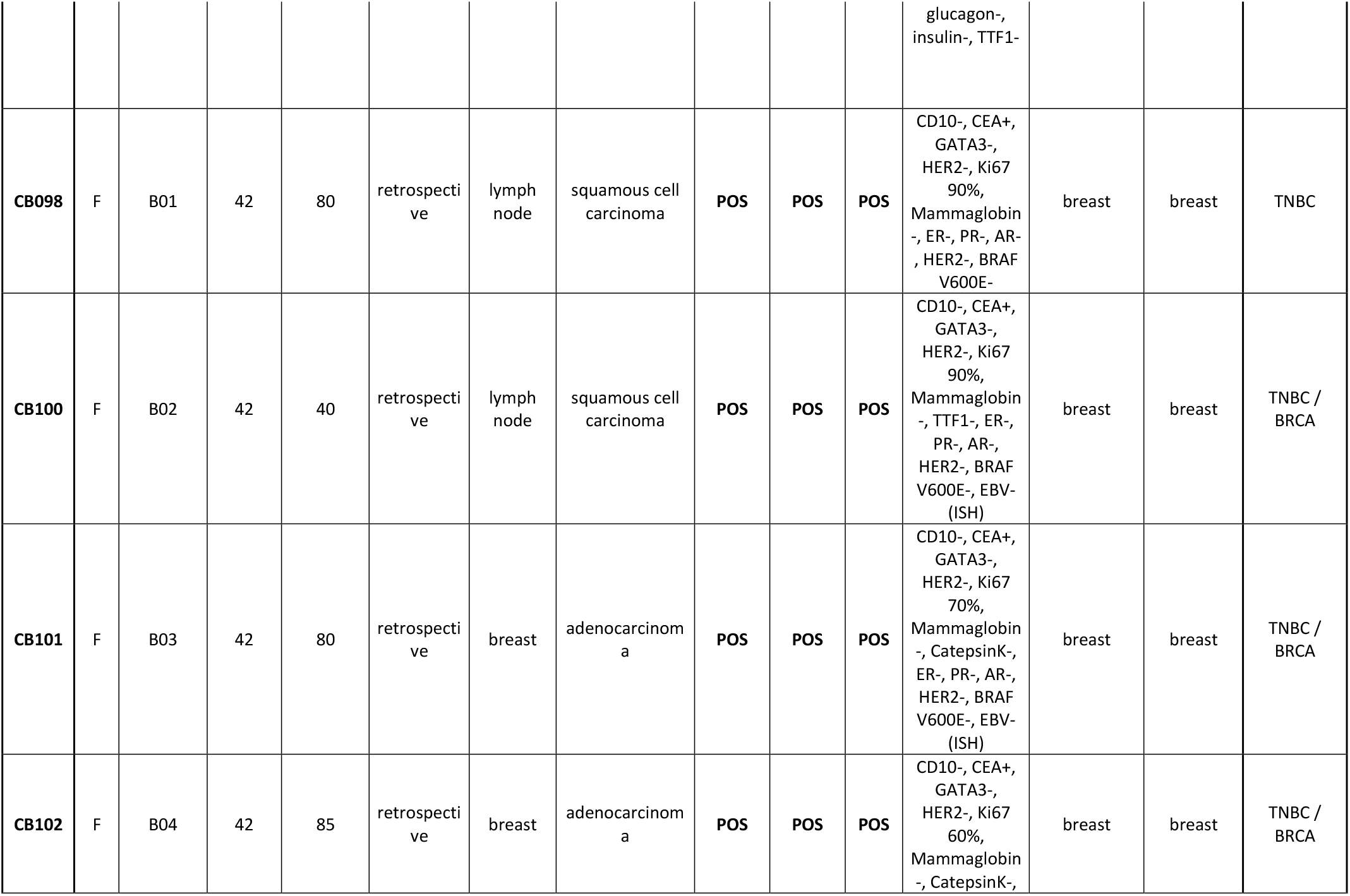
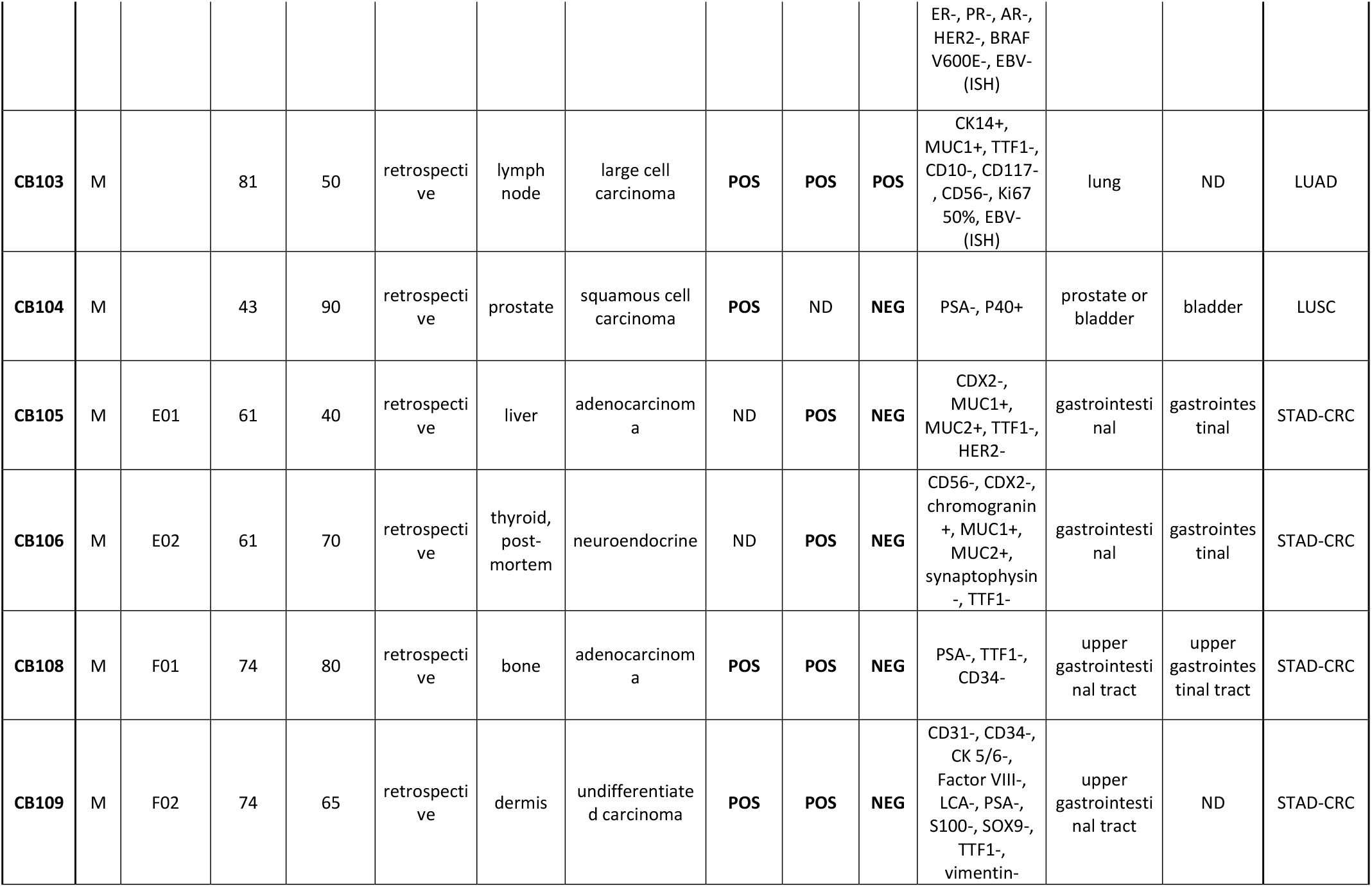
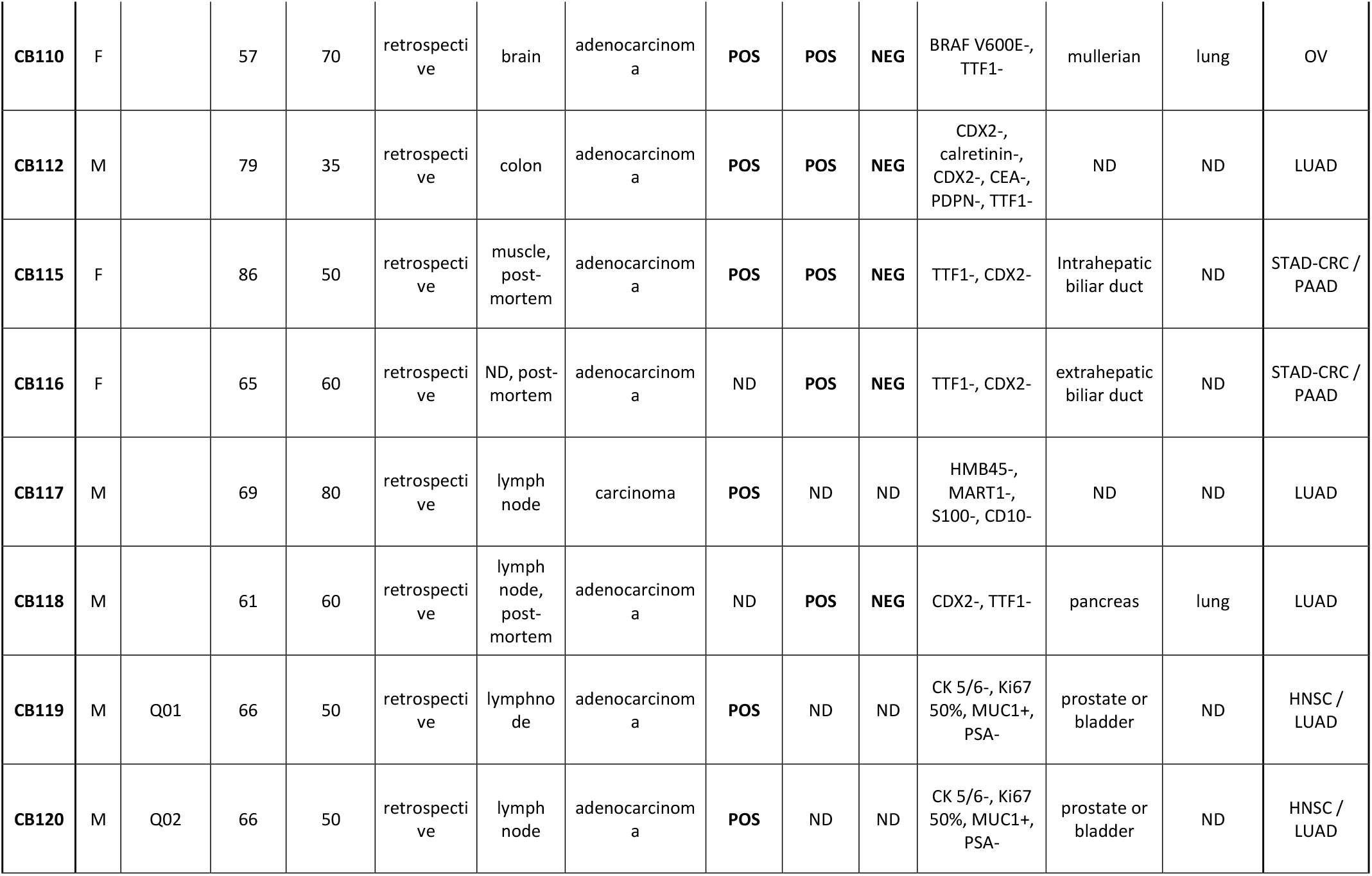
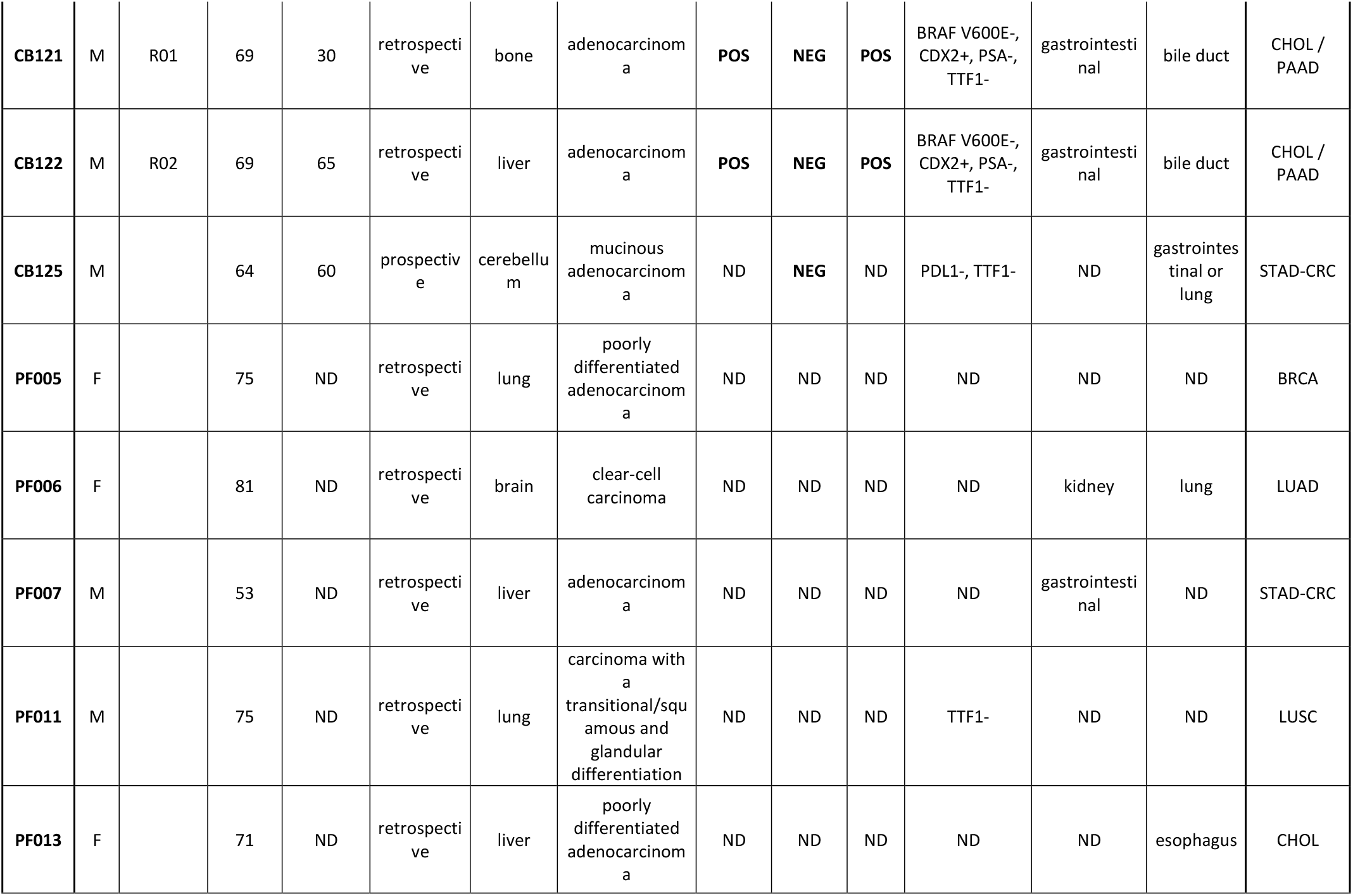
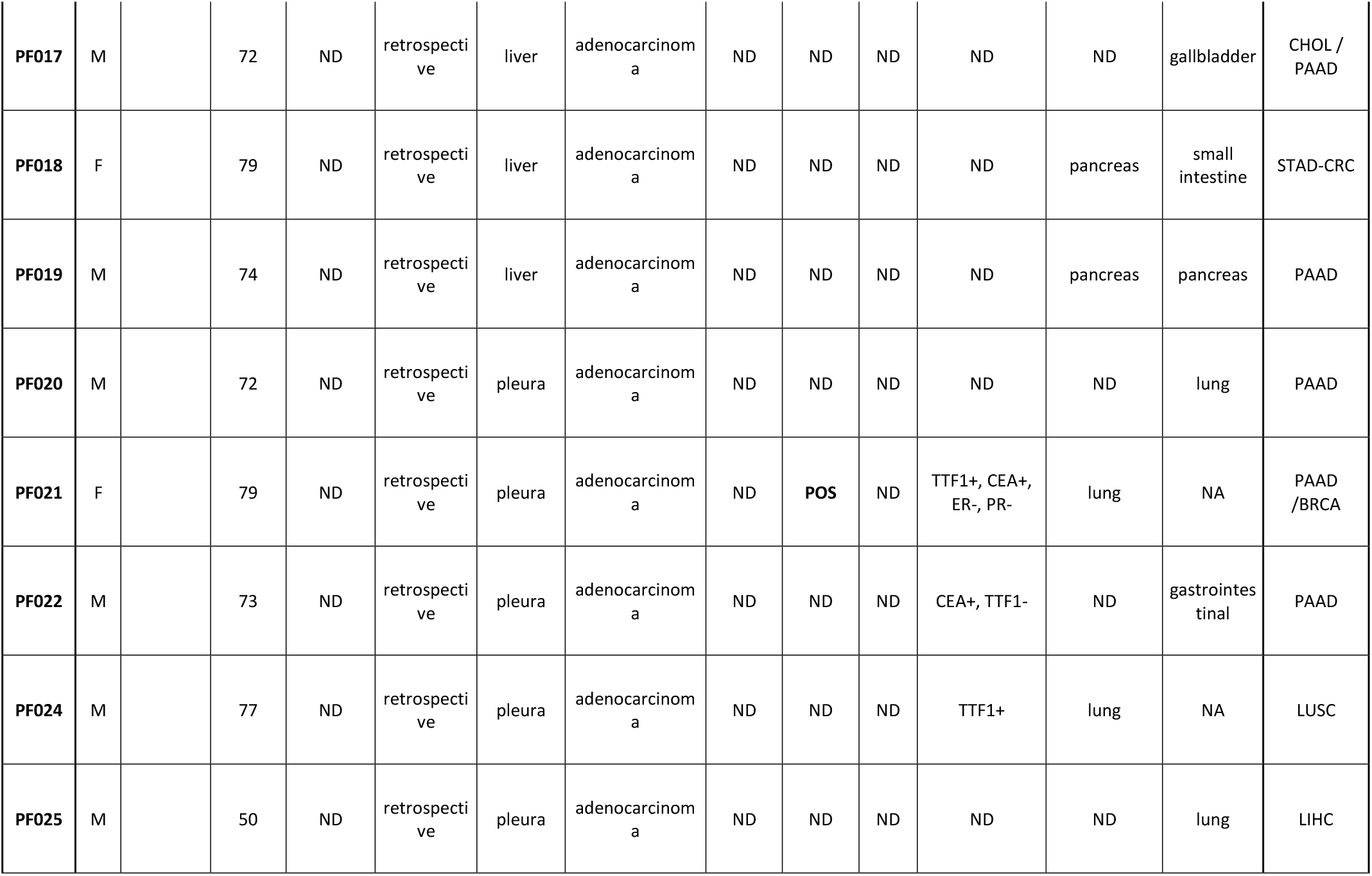
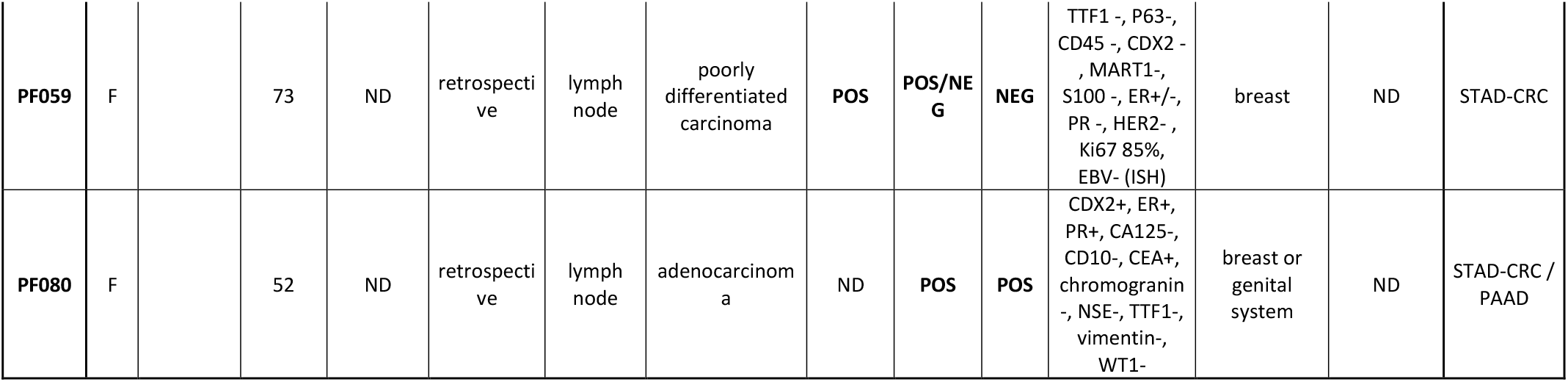
Prediction outcome in cancer of unknown primary site

| Sample ID | Sex | Multiple metastases | Age at diagnosis | Tumor cellularity (%) | Status | Biopsy site | Histotype | K WS | K7 | K20 | Other IHC testing | Pathological hypothesis | Clinical hypothesis | Molecular prediction (most probable primary tumor) |
| --- | --- | --- | --- | --- | --- | --- | --- | --- | --- | --- | --- | --- | --- | --- |
| CB002 | F |  | 65 | 50 | retrospective | liver | adenocarcinoma | ND | POS | NEG | CA125+, chromogranin-, ER-, GATA3- | ND | ND | STAD-CRC / BRCA |
| CB003 | F |  | 81 | 60 | retrospective | lymph node | carcinoma | POS | ND | ND | S100-, CD10-, TTF1-, ER-, PR-, HMB45-, GATA3- | ND | ND | HNSC / LUSC |
| CB011 | F |  | 75 | 85 | retrospective | lymph node | carcinoma | POS | POS | ND | WT1+, vimentin +, chromogranin +/-, synaptophysin +/-, calretinin-, CD10-, ER-, PR-, p63-, CD45-, CDX2- | mullerian or kidney | ND | PAAD |
| CB012 | M |  | 76 | 30 | retrospective | bone marrow | adenocarcinoma | POS/NEG | NEG | NEG | S100+, PSA +/-, CK14-, CK18-, CK 19-, ER-, PR-, HMB45-, MUC1+, TTF1-, vimentin- | prostate | melanoma | CHOL / LIHC |
| CB013 | F |  | 71 | 70 | retrospective | lymph node | mucinous adenocarcinoma | POS | NEG/POS | NEG | ER-, PR-, TTF1-, CDX2- | gastrointestinal | ND | STAD-CRC / PAAD |
| **CB014** | F |  | 47 | 50 | retrospective | lymph node | papillary adenocarcinoma | **POS/NEG** | **POS/NEG** | **NEG** | vimentin +/-, actin-, CK14+, ER-, PR-, HER2-, TTF1-, thyroglobulin-, WT1-, GATA3+, P40-, PAX8- | TN breast or thyroid | ND | TNBC |
| **CB033** | F |  | 77 | 60 | prospective | lymph node | carcinoma | **POS** | **POS** | **NEG** | CK 5-6+, CK14+, GATA3+/-, ER-, PR-, AR, HER2-, P63+, P40+, synaptophysin-, PAX8-, napsin A-WT1-, TTF1- | sudoriparous gland | breast | BRCA |
| **CB053** | F |  | 60 | 75 | prospective | liver | adenocarcinoma | ND | **POS** | **NEG** | TTF1+, ALK-, KRAS G34T (pyrosequencing) | lung | ND | BRCA |
| **CB054** | F |  | 59 | 70 | prospective | lung | carcinoma | **POS** | ND | ND | GATA3+, ER-, PR-, HER2-, Ki67 96%, p40-, p63-, TTF1-, CDX2- | breast | ND | BRCA |
| **CB055** | F |  | 49 | 50 | prospective | lymph node | adenocarcinoma | **POS** | **POS** | **POS** | CDX2+, chromogranin-, synaptophysin-, CD56-, HER2-, MSI- | gastrointestinal | ND | CHOL |
| **CB061** | M |  | 60 | 70 | retrospective | kidney | adenocarcinoma | **POS** | **POS** | ND | AR+, CD10-, OCT4-, PSA-, RCC-, TTF1-, GATA3-, NKX34.1- | breast or kidney | ND | BRCA |
| **CB062** | M |  | 87 | 80 | prospectiv<br>e | seminal<br>vesicle | carcinoma | **POS** | **NEG** | **NEG** | CDX2+, SMA-,<br>CD34-,<br>desmin-,<br>HER2-,<br>Ki67:50%,<br>MART1-, MSI-,<br>NKX3.1-, S100- | gastrointesti<br>nal | ND | CHOL |
| **CB064** | M |  | 58 | 65 | prospectiv<br>e | liver | squamous<br>carcinoma | ND | **NEG** | **NEG** | P63+, CK14+,<br>chromogranin<br>-,<br>synaptophysin<br>-, CD45-, S100- | ND | ND | HNSC |
| **CB071** | F |  | 64 | 60 | prospectiv<br>e | liver | carcinoma | ND | **POS** | **NEG** | CEA +/-,<br>GATA3-, ER-,<br>PR-/+, TTF1-,<br>PAX8- | biliar duct | ND | CHOL |
| **CB090** | F |  | 63 | 30 | prospectiv<br>e | duodeno | adenocarcinom<br>a | ND | **POS** | **NEG** | CDX2 +/-,<br>MUC 1+, MUC<br>2-, MUC 5AC+,<br>MUC 6+/-,<br>HER2-, MSI-,<br>synaptophysin<br>-, TTF1- | gastrointesti<br>nal | ND | STAD-CRC |
| **CB095** | F |  | 70 | 80 | prospectiv<br>e | soft<br>tissue | squamous<br>carcinoma | ND | **POS** | **NEG** | CA125-,<br>CA15.3-, CDX2<br>+/-, desmin-,<br>ER-, GATA3-,<br>MUC1+, P40-,<br>PAX-8-, PD-L1<br>70%, ER-,<br>S100-,<br>synaptophysin<br>-, TTF1- | ND | ND | BLCA |
| **CB097** | F |  | 68 | 90 | retrospecti<br>ve | liver | neuroendocrine | ND | ND | ND | chromogranin<br>+,<br>synaptophysin<br>+, CDX2-,<br>gastrin-, glucagon-,<br>insulin-, TTF1- | gastrointesti<br>nal NET | liver | PAAD |
| **CB098** | F | B01 | 42 | 80 | retrospecti<br>ve | lymph<br>node | squamous cell<br>carcinoma | **POS** | **POS** | **POS** | CD10-, CEA+,<br>GATA3-,<br>HER2-, Ki67<br>90%,<br>Mammaglobin<br>-, ER-, PR-, AR-<br>, HER2-, BRAF<br>V600E- | breast | breast | TNBC |
| **CB100** | F | B02 | 42 | 40 | retrospecti<br>ve | lymph<br>node | squamous cell<br>carcinoma | **POS** | **POS** | **POS** | CD10-, CEA+,<br>GATA3-,<br>HER2-, Ki67<br>90%,<br>Mammaglobin<br>-, TTF1-, ER-,<br>PR-, AR-,<br>HER2-, BRAF<br>V600E-, EBV-<br>(ISH) | breast | breast | TNBC /<br>BRCA |
| **CB101** | F | B03 | 42 | 80 | retrospecti<br>ve | breast | adenocarcinom<br>a | **POS** | **POS** | **POS** | CD10-, CEA+,<br>GATA3-,<br>HER2-, Ki67<br>70%,<br>Mammaglobin<br>-, CatepsinK-,<br>ER-, PR-, AR-,<br>HER2-, BRAF<br>V600E-, EBV-<br>(ISH) | breast | breast | TNBC /<br>BRCA |
| **CB102** | F | B04 | 42 | 85 | retrospecti<br>ve | breast | adenocarcinom<br>a | **POS** | **POS** | **POS** | CD10-, CEA+,<br>GATA3-,<br>HER2-, Ki67<br>60%,<br>Mammaglobin<br>-, CatepsinK-, | breast | breast | TNBC /<br>BRCA |
|  |  |  |  |  |  |  |  |  |  |  | ER-, PR-, AR-,<br>HER2-, BRAF<br>V600E-, EBV-<br>(ISH) |  |  |  |
| **CB103** | M |  | 81 | 50 | retrospecti<br>ve | lymph<br>node | large cell<br>carcinoma | **POS** | **POS** | **POS** | CK14+,<br>MUC1+, TTF1-,<br>CD10-, CD117-<br>, CD56-, Ki67<br>50%, EBV-<br>(ISH) | lung | ND | LUAD |
| **CB104** | M |  | 43 | 90 | retrospecti<br>ve | prostate | squamous cell<br>carcinoma | **POS** | ND | **NEG** | PSA-, P40+ | prostate or<br>bladder | bladder | LUSC |
| **CB105** | M | E01 | 61 | 40 | retrospecti<br>ve | liver | adenocarcinom<br>a | ND | **POS** | **NEG** | CDX2-,<br>MUC1+,<br>MUC2+, TTF1-,<br>HER2- | gastrointesti<br>nal | gastrointes<br>tinal | STAD-CRC |
| **CB106** | M | E02 | 61 | 70 | retrospecti<br>ve | thyroid,<br>post-<br>mortem | neuroendocrine | ND | **POS** | **NEG** | CD56-, CDX2-,<br>chromogranin<br>+, MUC1+,<br>MUC2+,<br>synaptophysin<br>-, TTF1- | gastrointesti<br>nal | gastrointes<br>tinal | STAD-CRC |
| **CB108** | M | F01 | 74 | 80 | retrospecti<br>ve | bone | adenocarcinom<br>a | **POS** | **POS** | **NEG** | PSA-, TTF1-,<br>CD34- | upper<br>gastrointesti<br>nal tract | upper<br>gastrointes<br>tinal tract | STAD-CRC |
| **CB109** | M | F02 | 74 | 65 | retrospecti<br>ve | dermis | undifferentiate<br>d carcinoma | **POS** | **POS** | **NEG** | CD31-, CD34-,<br>CK 5/6-,<br>Factor VIII-,<br>LCA-, PSA-,<br>S100-, SOX9-,<br>TTF1-,<br>vimentin- | upper<br>gastrointesti<br>nal tract | ND | STAD-CRC |
| **CB110** | F |  | 57 | 70 | retrospecti<br>ve | brain | adenocarcinom<br>a | **POS** | **POS** | **NEG** | BRAF V600E-,<br>TTF1- | mullerian | lung | OV |
| **CB112** | M |  | 79 | 35 | retrospecti<br>ve | colon | adenocarcinom<br>a | **POS** | **POS** | **NEG** | CDX2-,<br>calretinin-,<br>CDX2-, CEA-,<br>PDPN-, TTF1- | ND | ND | LUAD |
| **CB115** | F |  | 86 | 50 | retrospecti<br>ve | muscle,<br>post-mortem | adenocarcinom<br>a | **POS** | **POS** | **NEG** | TTF1-, CDX2- | Intrahepatic<br>biliar duct | ND | STAD-CRC /<br>PAAD |
| **CB116** | F |  | 65 | 60 | retrospecti<br>ve | ND, post-mortem | adenocarcinom<br>a | ND | **POS** | **NEG** | TTF1-, CDX2- | extrahepatic<br>biliar duct | ND | STAD-CRC /<br>PAAD |
| **CB117** | M |  | 69 | 80 | retrospecti<br>ve | lymph<br>node | carcinoma | **POS** | ND | ND | HMB45-,<br>MART1-,<br>S100-, CD10- | ND | ND | LUAD |
| **CB118** | M |  | 61 | 60 | retrospecti<br>ve | lymph<br>node,<br>post-mortem | adenocarcinom<br>a | ND | **POS** | **NEG** | CDX2-, TTF1- | pancreas | lung | LUAD |
| **CB119** | M | Q01 | 66 | 50 | retrospecti<br>ve | lymphno<br>de | adenocarcinom<br>a | **POS** | ND | ND | CK 5/6-, Ki67<br>50%, MUC1+,<br>PSA- | prostate or<br>bladder | ND | HNSC /<br>LUAD |
| **CB120** | M | Q02 | 66 | 50 | retrospecti<br>ve | lymph<br>node | adenocarcinom<br>a | **POS** | ND | ND | CK 5/6-, Ki67<br>50%, MUC1+,<br>PSA- | prostate or<br>bladder | ND | HNSC /<br>LUAD |
| **CB121** | M | R01 | 69 | 30 | retrospecti<br>ve | bone | adenocarcinom<br>a | **POS** | **NEG** | **POS** | BRAF V600E-,<br>CDX2+, PSA-,<br>TTF1- | gastrointesti<br>nal | bile duct | CHOL /<br>PAAD |
| **CB122** | M | R02 | 69 | 65 | retrospecti<br>ve | liver | adenocarcinom<br>a | **POS** | **NEG** | **POS** | BRAF V600E-,<br>CDX2+, PSA-,<br>TTF1- | gastrointesti<br>nal | bile duct | CHOL /<br>PAAD |
| **CB125** | M |  | 64 | 60 | prospectiv<br>e | cerebellu<br>m | mucinous<br>adenocarcinom<br>a | ND | **NEG** | ND | PDL1-, TTF1- | ND | gastrointes<br>tinal or<br>lung | STAD-CRC |
| **PF005** | F |  | 75 | ND | retrospecti<br>ve | lung | poorly<br>differentiated<br>adenocarcinom<br>a | ND | ND | ND | ND | ND | ND | BRCA |
| **PF006** | F |  | 81 | ND | retrospecti<br>ve | brain | clear-cell<br>carcinoma | ND | ND | ND | ND | kidney | lung | LUAD |
| **PF007** | M |  | 53 | ND | retrospecti<br>ve | liver | adenocarcinom<br>a | ND | ND | ND | ND | gastrointesti<br>nal | ND | STAD-CRC |
| **PF011** | M |  | 75 | ND | retrospecti<br>ve | lung | carcinoma with<br>a<br>transitional/squ<br>amous and<br>glandular<br>differentiation | ND | ND | ND | TTF1- | ND | ND | LUSC |
| **PF013** | F |  | 71 | ND | retrospecti<br>ve | liver | poorly<br>differentiated<br>adenocarcinom<br>a | ND | ND | ND | ND | ND | esophagus | CHOL |
| **PF017** | M |  | 72 | ND | retrospecti<br>ve | liver | adenocarcinom<br>a | ND | ND | ND | ND | ND | gallbladder | CHOL /<br>PAAD |
| **PF018** | F |  | 79 | ND | retrospecti<br>ve | liver | adenocarcinom<br>a | ND | ND | ND | ND | pancreas | small<br>intestine | STAD-CRC |
| **PF019** | M |  | 74 | ND | retrospecti<br>ve | liver | adenocarcinom<br>a | ND | ND | ND | ND | pancreas | pancreas | PAAD |
| **PF020** | M |  | 72 | ND | retrospecti<br>ve | pleura | adenocarcinom<br>a | ND | ND | ND | ND | ND | lung | PAAD |
| **PF021** | F |  | 79 | ND | retrospecti<br>ve | pleura | adenocarcinom<br>a | ND | **POS** | ND | TTF1+, CEA+,<br>ER-, PR- | lung | NA | PAAD<br>/BRCA |
| **PF022** | M |  | 73 | ND | retrospecti<br>ve | pleura | adenocarcinom<br>a | ND | ND | ND | CEA+, TTF1- | ND | gastrointes<br>tinal | PAAD |
| **PF024** | M |  | 77 | ND | retrospecti<br>ve | pleura | adenocarcinom<br>a | ND | ND | ND | TTF1+ | lung | NA | LUSC |
| **PF025** | M |  | 50 | ND | retrospecti<br>ve | pleura | adenocarcinom<br>a | ND | ND | ND | ND | ND | lung | LIHC |
| **PF059** | F |  | 73 | ND | retrospecti<br>ve | lymph<br>node | poorly<br>differentiated<br>carcinoma | **POS** | **POS/NE**<br>**G** | **NEG** | TTF1 -, P63-,<br>CD45 -, CDX2 -<br>, MART1-,<br>S100 -, ER+/-,<br>PR -, HER2- ,<br>Ki67 85%,<br>EBV- (ISH) | breast | ND | STAD-CRC |
| **PF080** | F |  | 52 | ND | retrospecti<br>ve | lymph<br>node | adenocarcinom<br>a | ND | **POS** | **POS** | CDX2+, ER+,<br>PR+, CA125-,<br>CD10-, CEA+,<br>chromogranin<br>-, NSE-, TTF1-,<br>vimentin-,<br>WT1- | breast or<br>genital<br>system | ND | STAD-CRC /<br>PAAD |

### RNA extraction and cDNA conversion

Total RNA, including microRNAs, was isolated from the tumor FFPE sections using miRNeasy FFPE kit (Qiagen, Cat No. 217504; miRNeasy FFPE Handbook Qiagen, HB-0374-005). Deparaffinization was performed with xylene as described in *Appendix A*. Further instructions were followed from step 11 of the *Protocol: Purification of Total RNA, Including miRNA, from FFPE Tissue Sections*. RNA was eluted in 20-30 μL and frozen at −80°C. RNA yield and quality were assessed with NanoGenius Spectrophotometer (ONDA Spectrophotometer). All samples were suitable for the molecular testing.

RNA conversion to cDNA was performed using the miRCURY LNA RT Kit (Qiagen, Cat No. 339340; miRCURY LNA miRNA PCR Handbook, HB-2431-002). The 10 μL-reaction mix was prepared for each sample mixing: 2 μL of 5x Reaction Buffer, 4.5 μL of nuclease free water, 1 μL of enzyme mix, 0.5 μL of UniSp6 RNA spike-in and 2 μL of diluted RNA (10 ng of total RNA). The resulting cDNA was stored in LoBind DNA Eppendorf tubes (Eppendorf, 0030108051) at −20°C. For each sample, a RT-qPCR was performed as quality control step using miRCURY LNA miRNA PCR Assays (Qiagen) to test UniSp6 (Cat No. YP00203954) and SNORD44 (Cat No. YP00203902) targets. UniSp6 threshold cycle (Ct) informs about the RT reaction efficiency. SNORD44 was tested to assess RNA integrity and amplifiability and to establish the cDNA dilution prior to digital droplet PCR (ddPCR) analysis. For SNORD44 Ct ranging 24-30 (threshold set at 160), cDNA was diluted 1:50; for Ct below 24, cDNA was diluted 1:100-1:200 and when Ct was higher than 30, the RT was repeated again using undiluted RNA and qPCR analysis repeated. cDNA was further diluted 1:10 in miR-21-5p and UniSP6 wells. Applying these criteria, we prevented ddPCR saturation problems or low miRNA expression levels in ddPCR analysis.

### Droplet Digital PCR and data analysis

Pre-spotted custom plates (96-well format) were designed to comprehend 89 different miRCURY LNA miRNA primers (Qiagen), three assays for small nuclear or nucleolar RNAs as reference candidates (SNORD44, SNORD48 and snRNAU6), two interplate calibrator assays (UniSp3), a control plate assay (UniSP6) and a no template control (NTC) as described in [34] (miRNA list in **Supplementary Table 2**).

EvaGreen-based Droplet digital PCR was performed as described in [35, 36, 34]. Thermal cycling conditions were: 95 °C for 5 min, then 40 cycles of 95 °C for 30 s and 58 °C for 1 min (ramping rate reduced to 2%), and three final steps at 4 °C for 5 min, 90 °C for 5 min and a 4 °C infinite hold. Droplet selection was performed individually for each well using QuantaSoft software v 1.7 (Bio-Rad). Final miRNA amounts (copies/ul) were obtained and normalized on 50^th^ percentile expression using GX v.14.9.1 software (Agilent Technologies).

### Tissue-of-origin prediction

Primary tumors (N=96) were used as training set as previously described [17]. Nearest shrunken centroids (NSC) algorithm [37] using the Prediction Analysis of Microarray for R (PAMR) tool [37] and the Least Absolute Shrinkage and Selection Operator (LASSO) model [38] were used to build up the classifiers. The PAM threshold was set to 0 leading to a classifier based on 88 miRNAs while the LASSO threshold was set to 0.019 leading to a classifier based on 48 miRNAs. Then, these classifiers were used to predict known and unknown/uncertain metastases tissue-of-origin. Both predictive models assign to every metastatic tumor a probability to be originated from each primary site. The variable gender was also taken into account to exclude not compatible molecular predictions (TGSC/PRAD in females and OV/UCEC in males). Results were compared with the indications of a possible primary site suggested by standard diagnostic workup and clinico-pathological assessment. Bootstrap approach (with N=100) was used to assess the performance (error rate) of the models in the training set.

### Cluster analysis

The hierarchical cluster analysis (with complete-linkage rule and Pearson Correlation distance) of primary tumor samples was obtained using GeneSpring GX v.14.9.1 software (Agilent Technologies) on ddPCR data normalized on the 50^th^ percentile.

### Survival analysis

Univariate survival analysis was performed using Kaplan-Meyer curves and the log rank test as implemented in survMisc R package. Overall survival (OS) was calculated considering the time lagging between diagnosis and death for any cause or the last follow-up. For each miRNA the optimal cut-off was estimated as the threshold on the ROC curves that maximize the sum of specificity and sensitivity in predicting CUP patients. Results were reported as *p* value, hazard ratio (HR) and 95% confidence intervals (CI). A *p* value ≤0.05 was considered significant.

## Results

### Multi-miRNA testing on archive samples with droplet digital PCR

We implemented a miRNA-signature for tumor primary site prediction integrating two published signatures [17, 39] plus 10 additional miRNAs (miR-661, miR-24-3p, miR-21-5p, miR-16-5p, miR-320a, miR-224-5p, miR-423-5p, miR-25-3p, miR-331-3p and miR-103a-3p) as detailed in **Supplementary Table 2**.

FFPE tissue is the most commonly available source of tumor material for molecular profiling in the clinical setting and miRNAs are extremely stable in FFPE blocks. Therefore, we developed an on-demand multi-miRNA assay capable of testing the absolute levels of 89 miRNAs in a 2-days timeframe compatible with standard diagnostic workup and with the amount of available material. The multi-miRNA assay is based on absolute miRNA quantification with EvaGreen Dye Droplet Digital PCR technology [34]. From a technical point of view, the assay provided good quality results for all tested archive FFPE samples. RNA was extracted from 2-5 slices of tumor FFPE blocks, then the tumor area was identified by experienced pathologists and macrodissected. An amount of 10 ng is sufficient to test all miRNAs in a single experiment, thus confirming the feasibility in a diagnostic setting.

We obtained the absolute copy number for all miRNAs included in our panel in the same Droplet Digital PCR experiment, with identical experimental conditions (annealing temperature and amount of primers), only adjusting the amount of input cDNA for miR-21-5p and UniSP6.

With the aim of establishing a reference set for cancer of unknown origin molecular profiling, we tested 96 primary tumors with our multi-miRNA assay, comprising 16 different tumor types and 19 histological classes, focusing on the most common CUP’s sites of origin identified at autopsy [40]. We obtained the expression matrix of the primary tumor dataset, constituted by tumors belonging to 19 different classes: LUAD, LUSC, PAAD, LIHC, CHOL, KIRC, KIRP, STAD, CRC, TGSC, OV, UCEC, BLCA, BRCA, TNBC, PRAD, SKCM, GI-NET and HNSC. An overview of the primary tumor samples for each histological subtype included in this study is reported in **Table 1**.

### Analysis of miRNA expression patterns

We evaluated the average levels of normalized expression of the 89-miRNA signature in the nineteen primary tumor types with cluster analysis (**Figure 1**). Each tumor type displays a peculiar pattern of miRNA expression, as expected. Nonetheless, we found some unexpected similarities and divergences among tumor types, which are worth mentioning. Specifically, miRNA expression of STAD and CRC was found to be consistently overlapping and partially intermixed with other gastrointestinal tumors (PAAD and GI-NET), as reported also in previous reports [17, 39, 41]. Due to this miRNA expression similarity, we decided to consider them as a single class (STAD-CRC) for molecular prediction. Similarly, kidney renal clear cell (KIRC) and papillary cell carcinomas (KIRP), showing similar miRNA expression patterns, were combined in the tumor class KICA. Tumors in female reproductive-system organs (OV and UCEC) were found to express similar yet distinct miRNA patterns as previously observed [17, 39, 41]. Moreover, lung cancers (both LUAD and LUSC) share a portion of their signatures with TNBC but not with other breast cancer subtypes (ER+, PR+, HER2+ tumors). TNBC shows a largely different pattern of miRNA expression when compared to other breast cancers, showing an unexpected similarity with HNSC instead. We could speculate that a common etiology associated to human papilloma virus (HPV) infection has been reported in both these tumor types [42–44]. Overall, this signature confirmed its potential in discriminating among 17 different tumor classes.

**Figure 1.**
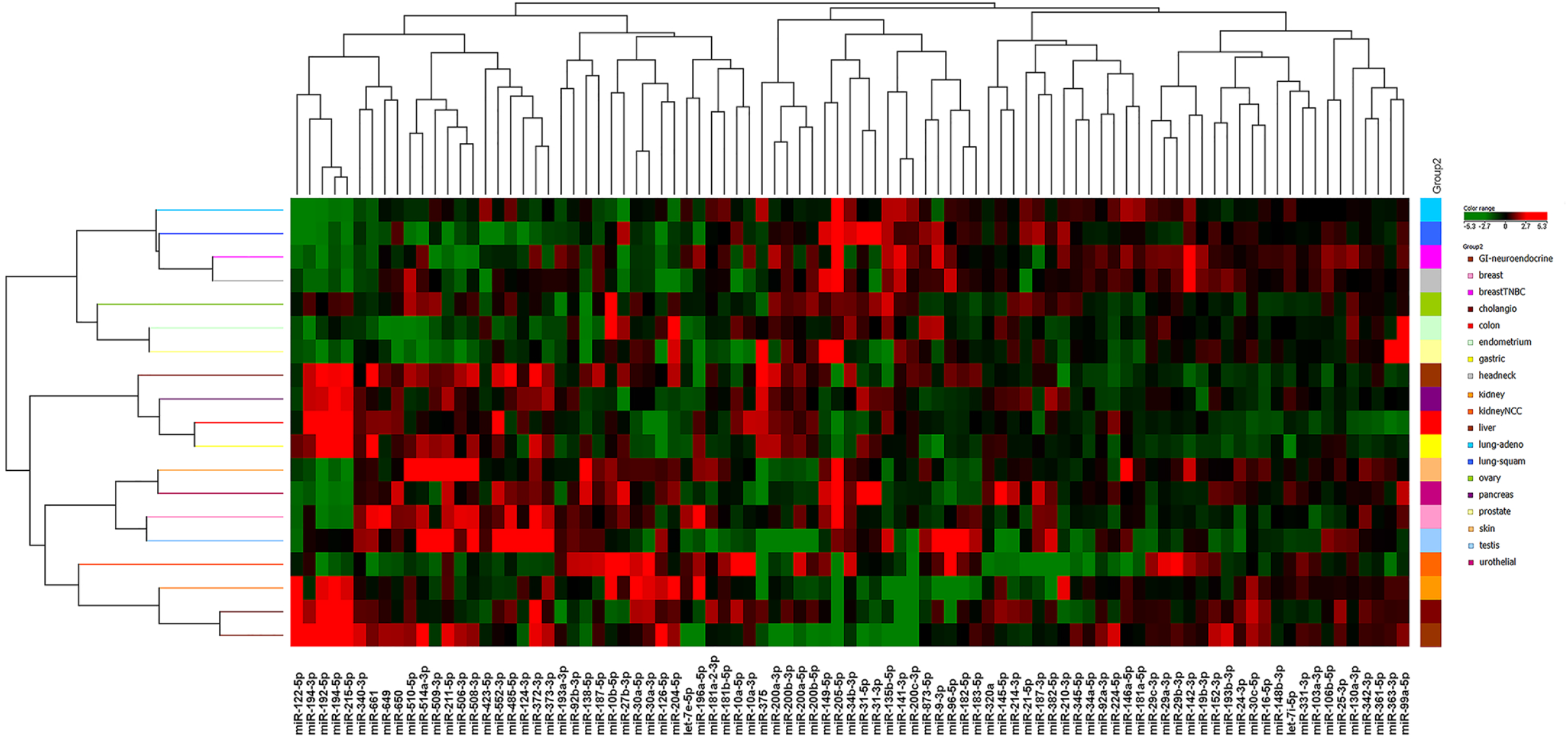
Cluster analysis of primary tumors. Heatmap representing 89-microRNA expression in nineteen different classes of primary carcinomas. Averaged, normalized miRNA levels in each tumor type were used for clustering analysis. Green indicates low expression, red indicates high expression.

### CUP predictive model generation

The final primary site prediction was performed using 88 out of 89 miRNA assays of our panel as miR-122-5p was excluded from the classifier due to its strong signal generated by the liver microenvironment in metastatic samples (**Supplementary Figure 1**).

We applied the Nearest Shrunken Centroids (NSC) using PAMR [37] and the Least Absolute Shrinkage and Selection Operator (LASSO) predictive models [38] developed by Tibshirani’s lab to our training set of primaries. To assess the performance of the predictive models on the training set we used a bootstrap approach. Error rates for each tumor class for both models are reported in **Supplementary Table 3**. Notably, the overall error rates for PAMR and LASSO were 34.3% and 35.1%, respectively. However, 11 of the 17 tumor classes (LIHC, LUSC, HNSC, BRCA, OV, CHOL, KICA, GI-NET, TGSC, STAD-CRC, SKCM, LUAD, UCEC, PRAD) had error rates significantly lower for both models (17.3% for PAMR and 22.7% for LASSO). Of note, PAMR seem to be considerably more accurate in the prediction of BRCA compared to LASSO, with no error reported; on the contrary, LASSO seem to be slightly more precise in the identification of UCEC, LUAD and SKCM. Both models had higher error rates in identifying correctly BLCA, PAAD, TNBC, HNSC, OV and CHOL classes; this might be explained by the low specificity of miRNA signatures for these primaries or by their small sample size. From these results it is clear that the two models behave similarly on some classes and complementarily in some others, thus we decided to take advantage of both classifiers for the second step of molecular prediction.

A small set of metastases of known origin (N=10) was assessed for molecular prediction (test set **Table 1**). Considering the two top predicted classes we obtained an accuracy of 80% for PAMR and of 60% for LASSO, as reported in **Supplementary Table 4**.

### CUP primary site prediction

Finally, both models were used to predict the primary site of 53 cancers of unknown/uncertain origin (CUPs). Given the tumor frequency, this is a remarkably large collection of cancers of unknown primary site whose histopathological and immunohistochemistry characteristics are detailed in **Supplementary Table 1** and **Table 2**. The prediction outcome is represented in **Figure 2** in which the top two primary sites predicted by both models for each CUP sample are reported. Using PAMR, the molecular prediction of 41 out of 53 CUPs (77.3%) predicted a primary site with a probability higher than 60%, half of them even higher than 90%. Using LASSO, the molecular prediction of 13 out of 53 CUPs (24.5%) predicted the first primary site with a probability higher than 60%, and 3 higher than 90%. We identified a subgroup of CUP patients for which it was very challenging to point out a tissue-of-origin using both models, thus suggesting an exceptionally undifferentiated phenotype. Of note, in the majority of cases LASSO and PAMR predictions were in agreement. Final molecular predictions are reported in **Table 2.** The most probable primary site which were selected based in the agreement between the two models and compatibility with the clinical/pathological information available. Considering the final predicted sites reported in **Table 2**, the most common tissues-of-origin were STAD-CRC (24.6%), PAAD (24.5%), BRCA (18.9%), LUAD or LUSC (15.1%), CHOL (15.1%), TNBC (8.2%) and HNSC (7.5%) and others at lower rates. Of note, no CUP was predicted to originate from TGSC or PRAD.

**Figure 2.**
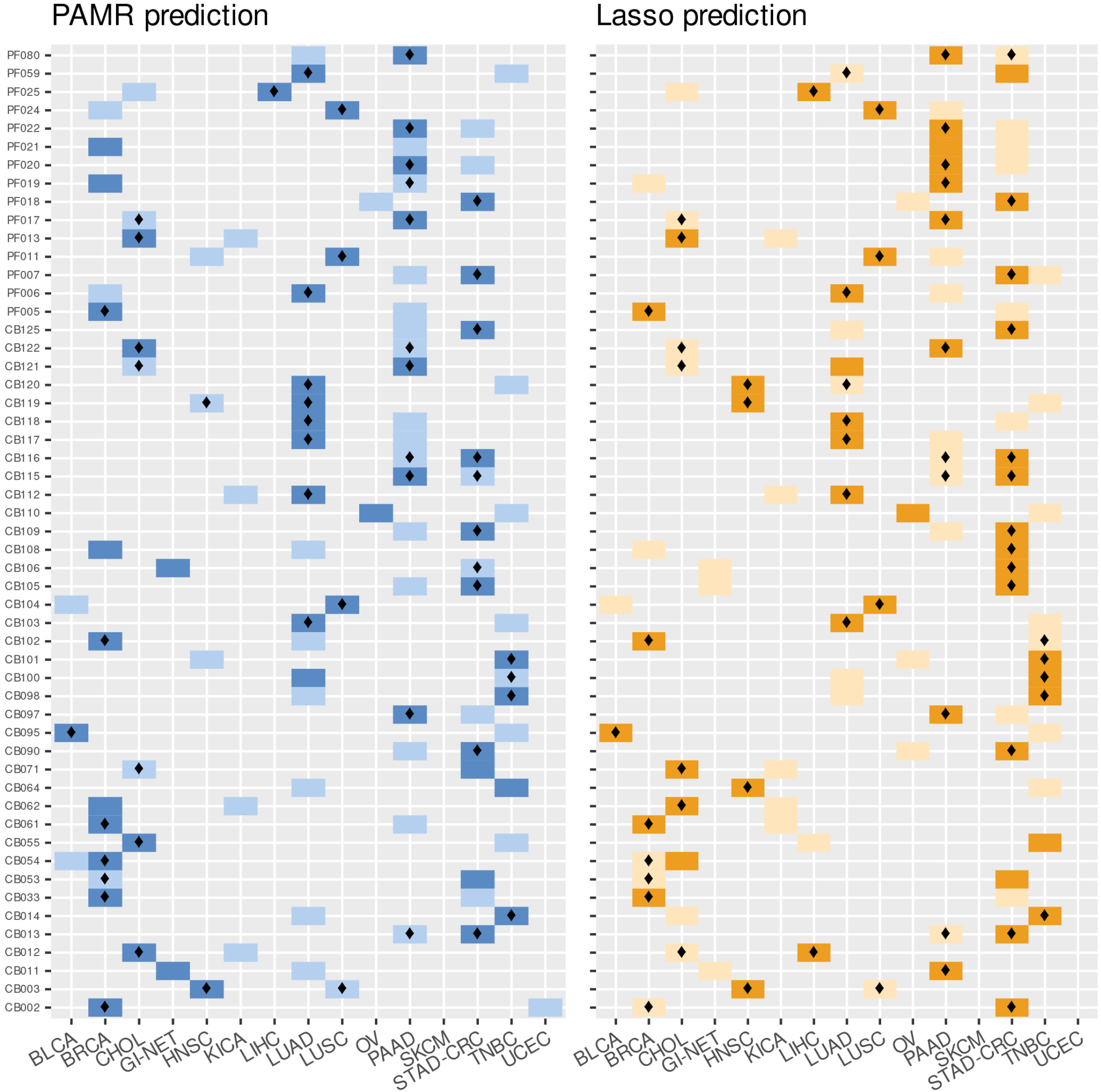
Prediction outcome of cancers of unknown origin using PAMR NSC and LASSO classifiers. For each of the 53 CUP sample (on the y-axis) the two top predicted primary tumors (x axis) are highlighted. PAMR first and second molecular predictions are reported with dark and light-blue squares, respectively. LASSO first and second molecular predictions are reported in dark- and lightorange, respectively. A diamond in the cell indicates those tissues-of-origin that are consistent with pathological and/or clinical information.

Interestingly, from five CUP patients we obtained a number of samples (N=2-4) derived from multiple metastatic sites, which were all tested with our assay. These samples were used to evaluate the consistency of our prediction. Symbolic is the case of a patient (#B) with an initial diagnosis of cancer of unknown primary (later attributed to a breast origin) from which we obtained a total number of four samples (CB098, CB100, CB101, CB102). In particular, CB098 and CB100 were obtained from two lymph nodes resected in 2010, while CB101 and CB102 derived from an invasive ductal breast cancer identified two years later, which was recognized as the primary site. Both PAMR and LASSO agreed to predict it as a BRCA or TNBC (**Table 2**). However, CB100 was predicted as LUAD (first) or TNBC (second) by PAMR classifier (**Figure 2**), probably due to the lower tumor cell fraction in this sample and the reported similarity in miRNA expression between breast and lung cancers [17]. Molecular predictions for the multiple metastases of the other patients (#E, #F, #Q and #R) reported concordant results for both models, in agreement with clinical or pathological hypotheses. Specifically, for #E (CB105 and CB106) and #F (CB108 and CB109) both models agreed to predict a gastrointestinal origin (STAD-CRC), which was also the first clinicopathological hypothesis. CB108 from #F patient had a different indication as most probable tissue-of-origin with PAMR classifier (breast); however, being derived from the bone it is probable that the sample had a compromised integrity. Molecular prediction for #R (CB121 and CB122) pointed out to a biliopancreatic origin, while for #Q (CB119 and CB120) the two metastatic samples were predicted to have the same origin, which was in this case lung or head and neck. CUP prediction probabilities with PAMR and LASSO models are reported in **Supplementary Table 5**.

### Association of microRNAs with CUP patients’ overall survival

We tested the performance of our 89-miRNA panel as prognostic test for CUP patients. Survival information was available for 34 CUP patients included in this study. We performed a survival analysis to test the association of miRNA expression with overall survival (**Supplementary Table 6**) finding 14 miRNAs with significant prognostic effect on CUP patients’ OS (**Table 3 and Figure 3**). The association between survival probability and miRNA expression was negative for 6 miRNAs (HR > 1) and positive for 8 miRNAs (HR < 1).

**Figure 3.**
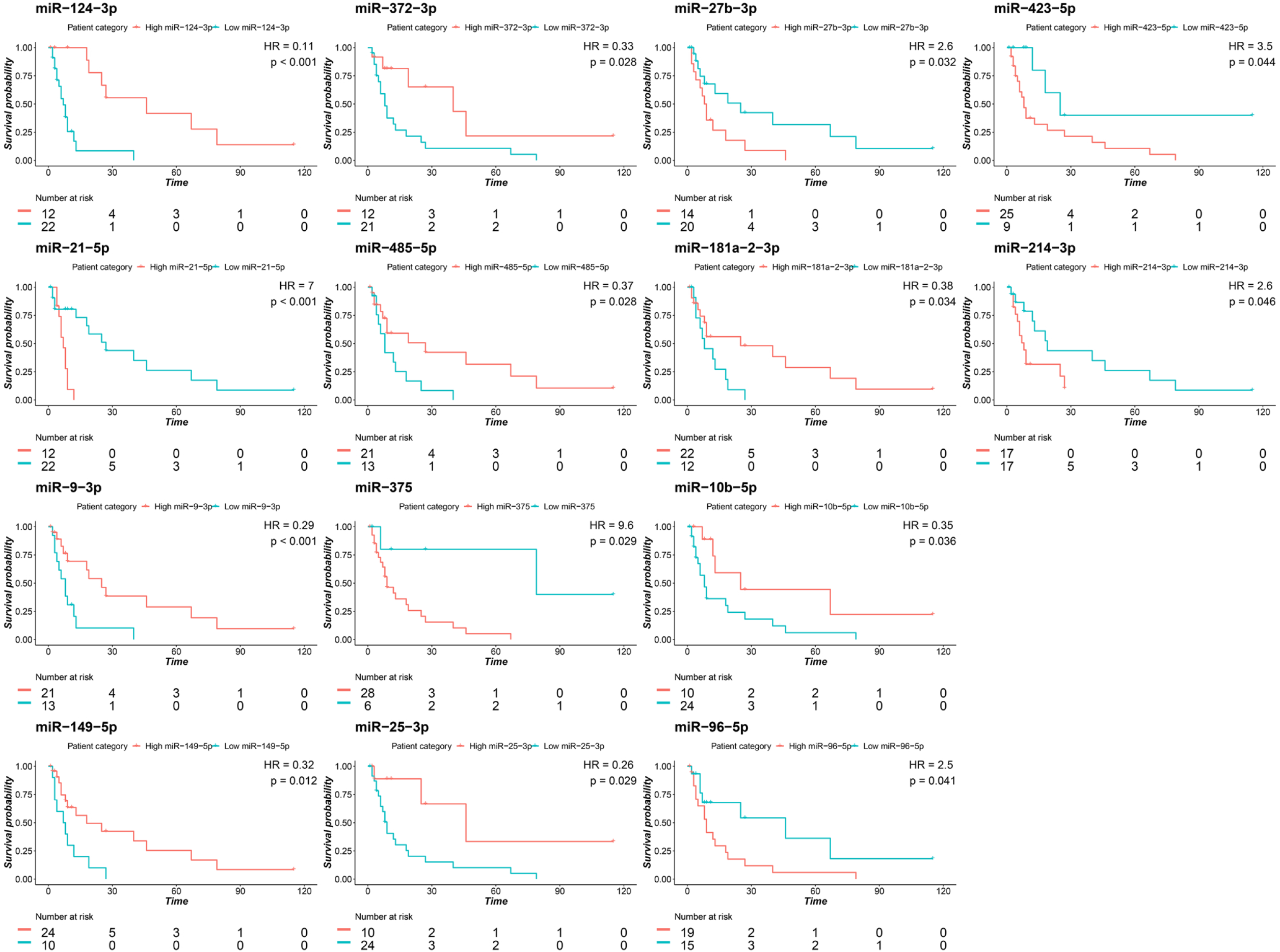
Kaplan–Meier OS curves based on the expression of 14 miRNAs in CUP patients. Survival plots showing significantly different OS curves in high- and low-miRNA expression cases. The threshold for each miRNA was established based on the best performing value at ROC analysis. A higher expression of 6 miRNAs is associated with shorter CUP survival and of 8 miRNAs with prolonged survival.

**Table 3.**
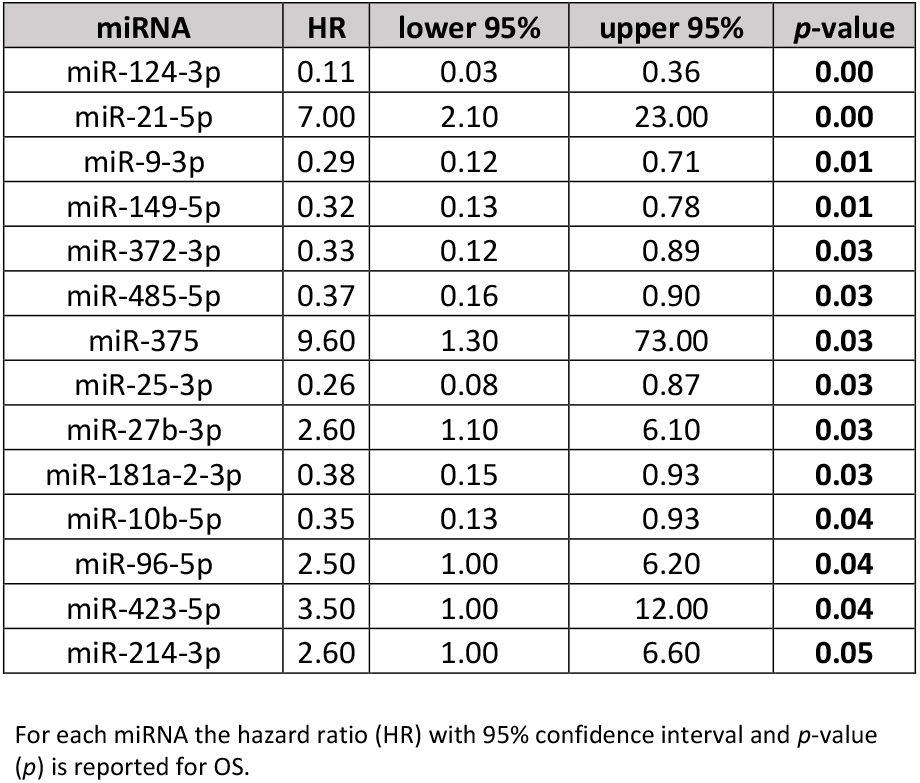
Association of miRNA expression with overall survival (significant miRNAs).

In particular, the miRNAs whose higher expression is associated with worse prognosis are miR-21-5p (*p*= 0.001), miR-375 (*p*=0.03), miR-27b-3p (*p*=0.03), miR-96-5p (*p*= 0.04), miR-423-5p (*p*=0.04) and miR-214-3p (*p*= 0.05). On the contrary, 8 miRNAs are positively associated with a prolonged survival: miR-124-3p (*p*= 0.0002), miR-9-3p (*p*= 0.01), miR-149-5p (*p*= 0.01), miR-372-3p (*p*= 0.03), miR-485-5p (*p*= 0.03), miR-25-3p (*p*= 0.03), miR-181a-2-3p (*p*= 0.03) and miR-10b-5p (*p*= 0.04).

## Discussion

The identification of the tissue-of-origin in metastatic cancers strongly relies on clinical information and histology as well as immunohistochemical evaluations but this diagnostic workup is sometimes ineffective and a fraction of primaries remains unidentified. Epitome of this scenario is metastatic cancer of unknown primary site (CUP), which presents by definition as an advanced cancer whose site of origin is not detectable nor presumable, despite an intensive clinical and pathological diagnostic workup [1]. CUPs represent an enigma both at the biological and pathological level, and an important under-researched clinical problem.

In the past decade, several molecular tests based on gene expression (GEP), microRNA or DNA methylation profile, were developed to improve primary site identification in cancers of unknown/uncertain origin. The underlying premise for these molecular profiling assays (reviewed in [45] and *Laprovitera at al*. (*Cancers, accepted*)) is that metastatic tumors preserve specific molecular signatures that match their primary site and can be used to identify their site-of-origin. Overall, these methods reach a prediction accuracy that ranges from 80 to 95% and have the potential to improve the diagnostic workup of CUP patients and guarantee the access to more therapeutic options. Indeed, NCCN occult primary guidelines recently assessed CUP molecular profiling as a potential provider of clinical benefit for patients. At the present time, CUP molecular profiling’s clinical utility need to be determined on a case-by-case basis, and clinical validation in large randomized phase III trials is still missing.

In this study, we developed a molecular assay to assess the expression of 89 miRNAs in tumor FFPE samples by using Droplet Digital PCR (ddPCR) and infer CUP primary site [34]. Our miRNA panel was determined merging two cancer-specific miRNA signatures previously identified in two microarray-based studies [17, 39]. To prevent the costs of large-scale technologies such as microarrays or sequencing, we opted for a focused number of selected miRNAs and the use of ddPCR technology. This assay allows the on-demand quantification of a focused panel of miRNAs per sample, at an affordable cost and in a 2-days timeframe. Droplet digital PCR technology provides miRNA absolute quantification without the requirement of standard curves, efficiency correction approaches or technical replicates typical of traditional quantitative PCR approaches [46]. In addition, EvaGreen-based ddPCR allows to precisely detect target miRNAs at levels down to 1 copy/μL [47].

As we hypothesized, an approach based on miRNA expression instead of gene expression profiles is very convenient since we were able to successfully analyze the totality of FFPE samples in our cohort (100% success rate), with no excluded sample due to technical issues.

In this study we analyzed the 89-miRNA profiles of 159 FFPE samples, including 53 CUPs, and successfully obtained a primary site prediction for all patients. We obtained a good prediction accuracy rate in metastatic cancers of known origin, and highly consistent results when assessing multiple metastases derived from the same CUP patient. These two settings provided an intrinsic validation of our combined predictive models.

As for CUP predictions, we observed consistency between our prediction outcomes and clinical and histo-pathological hypotheses, when they were available. In addition, we were able to successful analyze all 159 FFPE samples, with no excluded sample due to technical issues. The employment of two predictive models allowed us to obtain stronger results when both systems pointed out to the same tissue-of-origin. Of note, some CUPs were molecularly predicted as LUAD with a negativity for TTF1, which defines a subgroup of LUAD with unfavorable outcomes [48]. Our results provide further evidence of the translational potential of CUP molecular testing in general and miRNA testing in particular. With no intention to replace IHC testing, molecular assays can support the pathologists in narrowing the spectrum of possible primary sites of undifferentiated metastatic tumors. When no pathological hypothesis can be formulated, the miRNA-based molecular assay could aid the oncologists in their therapeutic choice, despite being necessary to demonstrate a benefit in a clinical setting.

The droplet digital PCR, miRNA-based assay herein applied has an accuracy comparable with other commercialized molecular profiling assays, but overcomes some limits of previous tools. Our molecular classifiers have the advantage to cover a wide variety of primary cancers, among the most likely to be CUP’s sites-of-origin; in particular, we can discriminate between 17 primary tumor subtypes. The ability to cover such number of tumor classes is an advantage if compared to other commercialized molecular assays e.g. the 10-genes qPCR assay (Veridex) that can classify only six different tumor types. Our prediction outcome on CUPs mostly overlap the frequency rates identified in post-mortem autopsy studies: lung (27%), pancreas (24%), liver or bile duct system (8%), kidney or adrenal (8%) or colon (7%) [40].

Three molecular assays were recently approved for CUP diagnostics in US: Pathwork Tissue of Origin Test (Pathwork Diagnostics), Cancer-TYPE ID (bioTheranostics), and miRview mets^2^ (Rosetta Genomics). The first is a microarray-based system that assessing the gene expression profiles (GEP) of 2000 genes claim to distinguish up to 15 tumor types. CancerTYPE ID is another GEP-based assay which evaluates by RT-qPCR the expression of a 92-gene signature and identifies the primary origin of up to 30 tumor types. Finally, miRview mets^2^ system, assessing the expression of 64 miRNAs by RT-qPCR, is able to distinguish up to 26 tumor types.

However, these assays included primary tumors that have little or no connection with CUPs. Our molecular tool is able to cover a high number of tumor classes, selected as the most common CUP tissues of origin. Our assay has a 100% success rate and requires a 2-days working time, which is compatible with a standard diagnostic workup and consistently shorter compared to other commercial assays that present a turnaround time of 5-11 days.

In addition to being faster, targeted and cost-effective in primary site identification; our assay could be easily combined with the analysis of druggable alterations, to select CUP therapy. However, further prospective clinical studies are necessary to evaluate their use in the clinics and to demonstrate its possible impact on CUP patients’ survival.

In conclusion, our study demonstrated that digital miRNA expression profiling of CUP samples has the potential to be employed in a clinical setting in FFPE tissue. Our molecular analysis can be performed on-request, concomitantly with the standard diagnostic workup and in association with genetic profiling, to offer valuable indication about the possible primary site thereby supporting treatment decisions.

## Declarations

### Ethics approval and consent to participate

The study was conducted in accordance with the Declaration of Helsinki, and the protocol was approved by the Ethics Committee Center Emilia-Romagna Region – Italy (protocol 130/2016/U/Tess) and Medical University of Graz (vote no. 30-520 ex 17/18). Prospective patients provided written informed consent.

### Consent for publication

All authors of the manuscript have read and agreed to its content.

### Availability of data and material

Droplet digital PCR data are available upon request.

### Authors’ contributions

Conception and design: M. Ferracin

Development of methodology: N. Laprovitera, M. Riefolo, E. Porcellini, M. Ferracin, C. Romualdi Acquisition of data: I. Garajova, F. Vasuri, A. Aigelsreiter, N. Dandachi, I. Berindan-Neagoe, A. Ardizzoni, D. Trerè, A. D’Errico

Analysis and interpretation of data: N. Laprovitera, M. Riefolo, E. Porcellini, F. Agostinis, G. Durante, G. Benvenuto, A. D’Errico, M. Pichler, M. Ferracin, C. Romualdi

Writing, review, and/or revision of the manuscript: N. Laprovitera, M. Riefolo, M. Pichler, A. D’Errico, M. Ferracin

Study supervision: M. Ferracin

## Acknowledgments

MR was a fellow of Fondazione Famiglia Parmiani, Bologna (Italy). NL is supported by eDIMES Lab funds from Bologna University.

## Supplementary information

**Supplementary Figure 1 - Plot of miR-122-5p expression in primary and metastatic tumors.** Normalized miR-122-5p expression was evaluated in liver and bile duct primary tumors, known to express this miRNA at high levels, and in metastatic tumors of known/unknown origin whose biopsy was obtained from the liver tissue or other sites. Liver metastases of known/unknown origin show high levels of miR-122-5p if compared to those derived from other sites, which is due to the very abundant expression of miR-122 in liver cells and its release in the tumor microenvironment.

**Supplementary Table 1 – Clinic-pathological features of 159 samples.**

**Supplementary Table 2 – List of miRNA assays in the custom ddPCR plate.**

**Supplementary Table 3. Error rates of the PAMR and LASSO models for each tumor class.**

**Supplementary Table 4. Primary site prediction in metastases of known origin.**

**Supplementary Table 5. CUP probabilities with PAMR and LASSO classifier models**

**Supplementary Table 6. Association of miRNA expression with CUP overall survival (all miRNAs).**

