## Supplementary figures and images for "Molecular diagnosis and prognosis of cancers of unknown-primary (CUPs): progress from a microRNA-based droplet digital PCR assay"

### Supplementary Figure 1

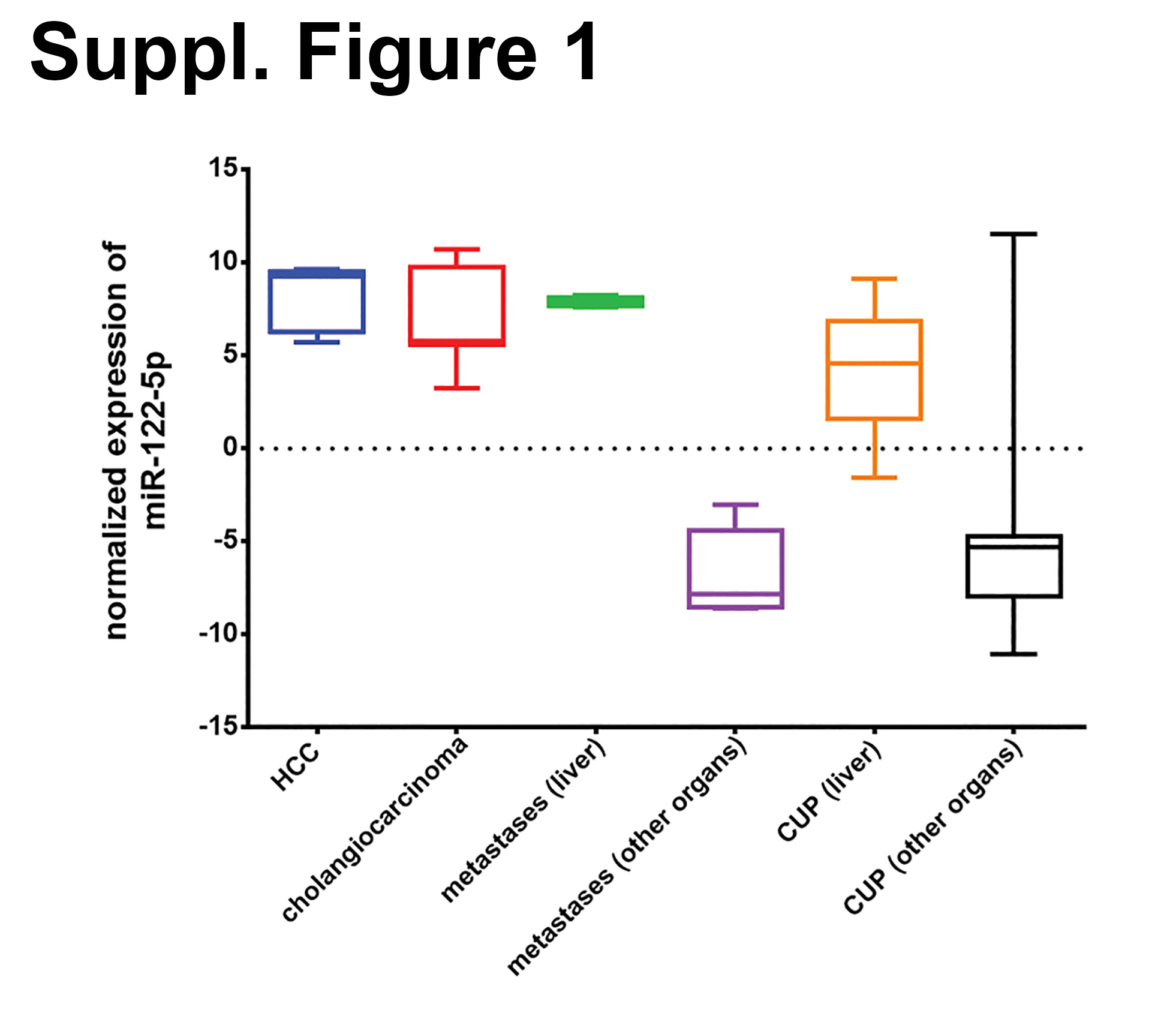
