## Supplementary Table 1 for "Molecular diagnosis and prognosis of cancers of unknown-primary (CUPs): progress from a microRNA-based droplet digital PCR assay"

**Supplementary table 1. Clinic-pathological features of 159 samples**

| Sample ID | Multiple metastases | Gender | Type | Age at diagnosis | Primary site | Sample site | Histology | FFPE tumor cellularity rate (%) | Stage/grading | Study |
|---|---|---|---|---|---|---|---|---|---|---|
| **BC001** | | ND | t | NA | breast | breast | lobular carcinoma | NA | NA | retrospective |
| **BC008** | | ND | t | NA | breast | breast | lobular carcinoma | NA | NA | retrospective |
| **BQ024** | | F | t | NA | breast | breast | TNBC | NA | NA | retrospective |
| **BQ033** | | F | t | NA | breast | breast | TNBC | NA | NA | retrospective |
| **BQ038** | | F | t | NA | breast | breast | TNBC | NA | NA | retrospective |
| **CB001** | | M | t | 75 | liver | liver | cholangiocarcinoma | NA | NA | retrospective |
| **CB002** | | F | CUP | 65 | NA | liver | adenocarcinoma | 50 | M1 | retrospective |
| **CB003** | | F | CUP | 81 | NA | soft tissue | muciparic adenocarcinoma | 60 | M1 | retrospective |
| **CB004** | | M | t | NA | testis | testis | germ cell seminomatous carcinoma | NA | NA | retrospective |
| **CB005** | | M | t | NA | testis | testis | germ cell seminomatous carcinoma | NA | NA | retrospective |
| **CB006** | | F | t | NA | pancreas | pancreas | adenocarcinoma | NA | NA | retrospective |
| **CB011** | | F | CUP | 75 | NA | lymph node | poorly differentiated carcinoma | 85 | M1 | retrospective |
| **CB012** | | M | CUP | 76 | NA | bone marrow | adenocarcinoma | 20 | M1 | retrospective |
| **CB013** | | F | CUP | 71 | NA | lymph node | mucinous adenocarcinoma | 70 | M1 | retrospective |
| **CB014** | | F | CUP | 47 | NA | lymph node | papillary adenocarcinoma | 50 | 3 | retrospective |
| **CB015** | | F | t | NA | lung | lung | adenocarcinoma | NA | NA | retrospective |
| **CB016** | | M | t | NA | lung | lung | adenocarcinoma | NA | NA | retrospective |
| **CB017** | | M | t | NA | lung | lung | adenocarcinoma | NA | NA | retrospective |
| **CB018** | | M | t | NA | lung | lung | adenocarcinoma | NA | NA | retrospective |
| **CB019** | | M | t | NA | gastrointestinal NET | gastrointestinal NET | gastrointestinal NET | NA | NA | retrospective |
| **CB020** | | F | t | NA | gastric | gastric | adenocarcinoma | NA | NA | retrospective |
| **CB021** | | F | t | NA | lung | lung | adenocarcinoma | NA | NA | retrospective |
| **CB022** | | F | t | NA | gastrointestinal NET | gastrointestinal NET | gastrointestinal NET | NA | NA | retrospective |
| **CB023** | | M | t | NA | gastric | gastric | adenocarcinoma | NA | NA | retrospective |
| **CB024** | | M | t | NA | gastrointestinal NET | gastrointestinal NET | gastrointestinal NET | NA | NA | retrospective |

| | | | | | | | | | | |
|---|---|---|---|---|---|---|---|---|---|---|
| **CB025** | | M | t | NA | gastrointestinal NET | gastrointestinal NET | gastrointestinal NET | NA | NA | retrospective |
| **CB026** | | M | t | NA | pancreas | pancreas | adenocarcinoma | NA | NA | retrospective |
| **CB028** | | M | t | NA | urothelial | urothelial | transitional cell carcinoma | NA | NA | retrospective |
| **CB032** | | M | t | NA | prostate | prostate | adenocarcinoma | NA | NA | retrospective |
| **CB033** | | F | CUP | 77 | NA | lymph node | carcinoma | 60 | NA | prospective |
| **CB034** | | M | t | NA | prostate | prostate | adenocarcinoma | NA | NA | retrospective |
| **CB035** | | M | t | NA | pancreas | pancreas | adenocarcinoma | NA | NA | retrospective |
| **CB036** | | M | t | NA | gastrointestinal NET | gastrointestinal NET | gastrointestinal NET | NA | NA | retrospective |
| **CB037** | | M | t | NA | kidney | kidney | clear-cell carcinoma | NA | NA | retrospective |
| **CB038** | | M | t | NA | kidney | kidney | clear-cell carcinoma | NA | NA | retrospective |
| **CB039** | | F | t | NA | kidney | kidney | clear-cell carcinoma | NA | NA | retrospective |
| **CB041** | | F | t | NA | urothelial | urothelial | transitional cell carcinoma | NA | NA | retrospective |
| **CB042** | | F | t | NA | urothelial | urothelial | transitional cell carcinoma | NA | NA | retrospective |
| **CB043** | | M | t | NA | urothelial | urothelial | transitional cell carcinoma | NA | NA | retrospective |
| **CB044** | | F | m | NA | colon/rectum | liver | Hepatic metastasis from colon adenocarcinoma | NA | NA | retrospective |
| **CB048** | | F | m | NA | breast | liver | Hepatic metastasis from breast | NA | NA | retrospective |
| **CB049** | | M | m | NA | pancreas | liver | Hepatic metastasis from pancreas | NA | NA | retrospective |
| **CB053** | | F | CUP | 60 | NA | liver | poorly differentiated, solid and micropapillary adenocarcinoma | 75 | NA | prospective |
| **CB054** | | F | CUP | 59 | NA | breast | carcinoma | 70 | NA | prospective |
| **CB055** | | F | CUP | 49 | NA | lymph node | adenocarcinoma | 50 | NA | prospective |
| **CB056** | | ND | t | NA | bile duct | bile duct | cholangiocarcinoma | NA | NA | retrospective |
| **CB057** | | ND | t | NA | bile duct | bile duct | cholangiocarcinoma | NA | NA | retrospective |
| **CB058** | | ND | t | NA | bile duct | bile duct | cholangiocarcinoma | NA | NA | retrospective |
| **CB059** | | ND | t | NA | bile duct | bile duct | cholangiocarcinoma | NA | NA | retrospective |
| **CB060** | | ND | t | NA | bile duct | bile duct | cholangiocarcinoma | NA | NA | retrospective |
| **CB061** | | M | CUP | 60 | NA | kidney | poorly differentiated adenocarcinoma | 70 | NA | retrospective |

| | | | | | | | | | | |
|---|---|---|---|---|---|---|---|---|---|---|
| **CB062** | | M | CUP | 87 | NA | prostate | carcinoma with a myxoid stroma and ring-shaped aspects with a castone | 80 | NA | prospective |
| **CB064** | | M | CUP | 58 | NA | liver | squamous carcinoma | 65 | NA | prospective |
| **CB065** | | ND | t | NA | testis | testis | germ cell seminomatous carcinoma | NA | NA | retrospective |
| **CB066** | | ND | t | NA | liver | liver | hepatocellular carcinoma | NA | NA | retrospective |
| **CB068** | | M | t | NA | testis | testis | germ cell seminomatous carcinoma | NA | NA | retrospective |
| **CB069** | | ND | t | NA | lung | lung | squamous | NA | NA | retrospective |
| **CB070** | | ND | t | NA | lung | lung | squamous | NA | NA | retrospective |
| **CB071** | | F | CUP | 64 | NA | liver | adenocarcinoma | 60 | G2 | prospective |
| **CB072** | | F | t | NA | endometrium | endometrium | adenocarcinoma | NA | NA | retrospective |
| **CB073** | | ND | m | NA | Head & Neck-Oropharyngeal | lung | adenocarcinoma | NA | NA | retrospective |
| **CB074** | | ND | t | NA | kidney | kidney | papillary cell carcinoma | NA | NA | retrospective |
| **CB075** | | ND | t | NA | liver | liver | hepatocellular carcinoma | NA | NA | retrospective |
| **CB076** | | ND | t | NA | kidney | kidney | papillary cell carcinoma | NA | NA | retrospective |
| **CB077** | | ND | t | NA | liver | liver | hepatocellular carcinoma | NA | NA | retrospective |
| **CB078** | | ND | t | NA | kidney | kidney | papillary cell carcinoma | NA | NA | retrospective |
| **CB080** | | M | t | NA | Head & Neck | Parotid glands | adenocarcinoma | NA | NA | retrospective |
| **CB081** | | M | t | NA | Head & Neck | Parotid glands | adenoid cystic carcinoma | NA | NA | retrospective |
| **CB082** | | M | t | NA | Head & Neck | tongue | epidermoid carcinoma of base of tongue | NA | NA | retrospective |
| **CB083** | | M | t | NA | Head & Neck | Rhinopharynx | carcinoma | NA | NA | retrospective |
| **CB085** | | F | t | NA | Head & Neck | tongue | epidermoid carcinoma of base of tongue | NA | NA | retrospective |
| **CB088** | | M | t | NA | Head & Neck | Parotid glands | adenocarcinoma | NA | NA | retrospective |
| **CB090** | | F | CUP | 63 | NA | duodeno (ampulla of Vater) | poorly differentiated carcinoma | 30 | G3 | prospective |
| **CB091** | | F | t | NA | ovary | ovary | ovarian serous carcinoma | NA | NA | retrospective |
| **CB093** | | F | t | NA | ovary | ovary | ovarian serous carcinoma | NA | NA | retrospective |
| **CB094** | | F | t | NA | ovary | ovary | ovarian serous carcinoma | NA | NA | retrospective |
| **CB095** | | F | CUP | 70 | NA | soft tissue | squamous carcinoma with sarcomatoid aspect | 80 | NA | prospective |

| | | | | | | | | | | |
|---|---|---|---|---|---|---|---|---|---|---|
| **CB097** | | F | CUP | 68 | NA | liver | neuroendocrine | 90 | M1 | retrospective |
| **CB098** | B01 | F | CUP | 42 | NA | lymph node | squamous cell carcinoma | 80 | NA | retrospective |
| **CB100** | B02 | F | CUP | 42 | NA | lymph node | squamous cell carcinoma | 40 | NA | retrospective |
| **CB101** | B03 | F | CUP | 42 | NA | breast | adenocarcinoma | 80 | NA | retrospective |
| **CB102** | B04 | F | CUP | 42 | NA | breast | adenocarcinoma | 85 | NA | retrospective |
| **CB103** | | M | CUP | 81 | NA | lymphnode | large cell carcinoma | 50 | NA | retrospective |
| **CB104** | | M | CUP | 43 | NA | prostate | squamous cell carcinoma | 90 | NA | retrospective |
| **CB105** | E01 | M | CUP | 61 | NA | liver | adenocarcinoma | 40 | NA | retrospective |
| **CB106** | E02 | M | CUP | 61 | NA | thyroid, post-mortem | neuroendocrine | 70 | NA | retrospective |
| **CB108** | F01 | M | CUP | 74 | NA | bone | adenocarcinoma | 80 | M1 | retrospective |
| **CB109** | F02 | M | CUP | 74 | NA | dermis | undifferentiated carcinoma | 65 | M1 | retrospective |
| **CB110** | | F | CUP | 57 | NA | brain | adenocarcinoma | 70 | M1 | retrospective |
| **CB112** | | M | CUP | 79 | NA | colon | adenocarcinoma | 35 | M1 | retrospective |
| **CB115** | | F | CUP | 86 | NA | muscle, post-mortem | adenocarcinoma | 50 | M1 | retrospective |
| **CB116** | | F | CUP | 65 | NA | ND, post-mortem | adenocarcinoma | 60 | M1 | retrospective |
| **CB117** | | M | CUP | 69 | NA | lymphnode | large cell carcinoma | 80 | M1 | retrospective |
| **CB118** | | M | CUP | 61 | NA | lymphnode, post-mortem | adenocarcinoma | 60 | M1 | retrospective |
| **CB119** | Q01 | M | CUP | 66 | NA | lymphnode | adenocarcinoma | 50 | M1 | retrospective |
| **CB120** | Q02 | M | CUP | 66 | NA | lymph node | adenocarcinoma | 50 | M1 | retrospective |
| **CB121** | R01 | M | CUP | 69 | NA | bone | adenocarcinoma | 30 | M1 | retrospective |
| **CB122** | R02 | M | CUP | 69 | NA | liver | adenocarcinoma | 65 | M1 | retrospective |
| **CB125** | | M | CUP | 64 | NA | cerebellum | mucinous adenocarcinoma | 60 | NA | prospective |
| **CRC05** | | M | t | NA | colon | colon | adenocarcinoma | NA | T3N0 | retrospective |
| **CRC27** | | M | t | NA | colon | colon | adenocarcinoma | NA | T3N0 | retrospective |
| **M016** | | M | t | NA | skin/melanoma | skin/melanoma | malignant melanoma | NA | NA | retrospective |
| **M017** | | M | t | NA | skin/melanoma | skin/melanoma | malignant melanoma | NA | NA | retrospective |

| | | | | | | | | | | |
|---|---|---|---|---|---|---|---|---|---|---|
| **M028** | | M | t | NA | skin/melanoma | skin/melanoma | malignant melanoma | NA | NA | retrospective |
| **M040** | | M | t | NA | skin/melanoma | skin/melanoma | malignant melanoma | NA | NA | retrospective |
| **PF001** | | F | t | 79 | breast | breast | lobular carcinoma | NA | 2 | retrospective |
| **PF002** | | F | t | 59 | ovary | ovary | serous papillary carcinoma | NA | 3 | retrospective |
| **PF003** | | M | t | 71 | lung | lung | squamous cell carcinoma | NA | 3 | retrospective |
| **PF004** | | F | t | 44 | breast | breast | ductal carcinoma | NA | 3 | retrospective |
| **PF005** | | F | CUP | 75 | NA | lung | poorly differentiated adenocarcinoma | NA | NA | retrospective |
| **PF006** | | F | CUP | 81 | NA | brain | clear-cell carcinoma | NA | NA | retrospective |
| **PF007** | | M | CUP | 53 | NA | liver | adenocarcinoma | NA | NA | retrospective |
| **PF010** | | M | m | 62 | lung | brain | adenocarcinoma | NA | 4 | retrospective |
| **PF011** | | M | CUP | 75 | NA | lung | carcinoma with a transitional/squamous and glandular | NA | NA | retrospective |
| **PF013** | | F | CUP | 71 | NA | liver | Poorly differentiated adenocarcinoma | NA | NA | retrospective |
| **PF017** | | M | CUP | 72 | NA | liver | adenocarcinoma | NA | NA | retrospective |
| **PF018** | | F | CUP | 79 | NA | liver | adenocarcinoma | NA | NA | retrospective |
| **PF019** | | M | CUP | 74 | NA | liver | adenocarcinoma | NA | NA | retrospective |
| **PF020** | | M | CUP | 72 | NA | pleura | adenocarcinoma | NA | NA | retrospective |
| **PF021** | | F | CUP | 79 | NA | pleura | adenocarcinoma | NA | NA | retrospective |
| **PF022** | | M | CUP | 73 | NA | pleura | adenocarcinoma | NA | NA | retrospective |
| **PF024** | | M | CUP | 77 | NA | pleura | adenocarcinoma | NA | NA | retrospective |
| **PF025** | | M | CUP | 50 | NA | pleura | adenocarcinoma | NA | NA | retrospective |
| **PF026** | | M | t | 51 | colon | colon | adenocarcinoma | NA | 4 | retrospective |
| **PF027** | | F | t | 75 | kidney | kidney | clear-cell carcinoma | NA | 3 | retrospective |
| **PF028** | | F | t | 79 | gastric | gastric | adenocarcinoma | NA | 3 | retrospective |
| **PF029** | | M | t | 75 | gastric | gastric | adenocarcinoma | NA | 3 | retrospective |
| **PF031** | | M | t | 74 | liver | liver | hepatocellular carcinoma | NA | 2 | retrospective |
| **PF038** | | F | t | 71 | breast | breast | ductal and lobular carcinoma | NA | 2 | retrospective |

| | | | | | | | | | | |
|---|---|---|---|---|---|---|---|---|---|---|
| **PF043** | | M | t | 63 | colon/rectum | colon/rectum | adenocarcinoma | NA | 2 | retrospective |
| **PF045** | | M | t | 63 | colon | colon | adenocarcinoma | NA | 2 | retrospective |
| **PF051** | | M | t | 67 | lung | lung | adenosquamous carcinoma | NA | 3 | retrospective |
| **PF059** | | F | CUP | 73 | NA | lymph node | poorly differentiated carcinoma | NA | NA | retrospective |
| **PF060** | | M | t | 67 | colon | colon | adenocarcinoma | NA | 2 | retrospective |
| **PF062** | | M | t | 69 | colon | colon | adenocarcinoma | NA | 2; T2N2 | retrospective |
| **PF065** | | M | t | 57 | pancreas | pancreas | adenocarcinoma | NA | 3 | retrospective |
| **PF066** | | F | t | 62 | pancreas | pancreas | ductal carcinoma | NA | 2 | retrospective |
| **PF071** | | F | t | 62 | kidney | kidney | clear-cell carcinoma | NA | 2 | retrospective |
| **PF075** | | M | t | 56 | skin/melanoma | skin/melanoma | malignant melanoma | NA | NA | retrospective |
| **PF079** | | M | m | 79 | skin/melanoma | gastric | malignat melanoma | NA | 4 | retrospective |
| **PF080** | | F | CUP | 52 | NA | lymph node | adenocarcinoma | NA | NA | retrospective |
| **PF081** | | F | t | 46 | ovary | ovary | mucinous adenocarcinoma | NA | 3 | retrospective |
| **PF082** | | F | t | 44 | ovary | ovary | mucinous cystadenocarcinoma | NA | NA | retrospective |
| **PF085** | | M | t | 77 | gastric | gastric | adenocarcinoma | NA | 3 | retrospective |
| **PF086** | | F | m | 75 | gastric | liver | adenocarcinoma | NA | 3 | retrospective |
| **PF087** | | M | m | 86 | prostate | colon | adenocarcinoma | NA | NA | retrospective |
| **PF088** | | F | m | 60 | endometrium | skin/melanoma | adenocarcinoma | NA | NA | retrospective |
| **PF090** | | F | t | 59 | endometrium | endometrium | adenocarcinoma | NA | 2 | retrospective |
| **PF092** | | F | t | 65 | endometrium | endometrium | adenocarcinoma | NA | 2 | retrospective |
| **PF093** | | M | t | 69 | prostate | prostate | adenocarcinoma | NA | 3; pT3a-pNX | retrospective |
| **PF094** | | M | t | 69 | prostate | prostate | adenocarcinoma | NA | 3; pT3a-pNX | retrospective |
| **PF095** | | F | m | 65 | kidney | pericardium | clear-cell carcinoma | NA | 2 | retrospective |
| **PF100** | | M | t | 73 | prostate | prostate | adenocarcinoma | NA | 4; pT3b | retrospective |
| **PF104** | | F | t | 80 | endometrium | endometrium | adenocarcinoma | NA | 2 | retrospective |
| **PF109** | | F | t | 80 | endometrium | endometrium | adenocarcinoma | NA | 2 | retrospective |

| | | | | | | | | | | |
|---|---|---|---|---|---|---|---|---|---|---|
| **PF30A** | | F | t | 85 | liver | liver | hepatocellular carcinoma | NA | 1 | retrospective |
| **PF30B** | | F | t | 85 | liver | liver | hepatocellular carcinoma | NA | 1 | retrospective |
| **PF77A** | | M | t | 68 | skin/melanoma | skin/melanoma | malignant melanoma | NA | 4; pT3b | retrospective |
| **PF77B** | | M | t | 68 | skin/melanoma | skin/melanoma | malignant melanoma | NA | 4; pT3b | retrospective |

*t= primary tumor; m= metastasis with known origin; CUP= cancer of unknown primary site; NET= neuroendocrine tumor; NA= not available; TNBC= triple-negative breast cancer.*
