## Supplementary Table 2 for "Molecular diagnosis and prognosis of cancers of unknown-primary (CUPs): progress from a microRNA-based droplet digital PCR assay"

**Supplementary table 2. List of miRNA assays in the custom ddPCR plate**

| microRNA | Plate well | Target sequence | Previous nomenclature | Qiagen LNA assay | Reference |
|---|---|---|---|---|---|
| hsa-let-7e-5p | A01 | UGAGGUAGGAGGUUGUAUAGUU | hsa-let-7e | YP00205711 | {Rosenfeld, 2008 #21} |
| hsa-let-7i-5p | A02 | UGAGGUAGUAGUUUGUGCUGUU | hsa-let-7i | YP00204394 | {Rosenfeld, 2008 #21} |
| hsa-miR-10a-5p | A03 | UACCCUGUAGAUCCGAAUUUGUG | hsa-miR-10a | YP00204778 | {Ferracin, 2011 #133} |
| hsa-miR-10a-3p | A04 | CAAAUUCGUAUCUAGGGGAAUA | hsa-miR-10a* | YP00205688 | {Ferracin, 2011 #133} |
| hsa-miR-10b-5p | A05 | UACCCUGUAGAACCGAAUUUGUG | hsa-miR-10b | YP00205637 | {Rosenfeld, 2008 #21} |
| hsa-miR-106b-5p | A06 | UAAAGUGCUGACAGUGCAGAU | hsa-miR-106b | YP00205884 | {Rosenfeld, 2008 #21} |
| hsa-miR-122-5p | A07 | UGGAGUGUGACAAUGGUGUUUG | hsa-miR-122 | YP00205664 | {Ferracin, 2011 #133} |
| hsa-miR-124-3p | A08 | UAAGGCACGCGGUGAAUGCCAA | hsa-miR-124 | YP00206026 | {Rosenfeld, 2008 #21} |
| hsa-miR-126-5p | A09 | CAUUAUUACUUUUGGUACGCG | hsa-miR-126* | YP00206010 | {Ferracin, 2011 #133} |
| hsa-miR-130a-3p | A10 | CAGUGCAAUGUUAAAAGGGCAU | hsa-miR-130a | YP00204658 | {Rosenfeld, 2008 #21} |
| hsa-miR-135b-5p | A11 | UAUGGCUUUUCAUUCCUAUGUGA | hsa-miR-135b | YP00204130 | {Ferracin, 2011 #133} |
| UniSp3 IPC | A12 | | | YP02119288 | |
| hsa-miR-138-5p | B01 | AGCUGGUGUUGUGAAUCAGGCCG | hsa-miR-138 | YP00206078 | {Rosenfeld, 2008 #21} |
| hsa-miR-141-3p | B02 | UAACACUGUCUGGUAAAGAUGG | hsa-miR-141 | YP00204504 | {Ferracin, 2011 #133} |
| hsa-miR-142-3p | B03 | UGUAGUGUUUCCUACUUUAUGGA | hsa-miR-142-3p | YP00204291 | {Rosenfeld, 2008 #21} |
| hsa-miR-145-5p | B04 | GUCCAGUUUUCCCAGGAAUCCCU | hsa-miR-145 | YP00204483 | {Ferracin, 2011 #133} |
| hsa-miR-146a-5p | B05 | UGAGAACUGAAUUCCAUGGGUU | hsa-miR-146a | YP00204688 | {Ferracin, 2011 #133} |
| hsa-miR-148b-3p | B06 | UCAGUGCAUCACAGAACUUUGU | hsa-miR-148b | YP00203906 | {Rosenfeld, 2008 #21} |
| hsa-miR-149-5p | B07 | UCUGGCUCCGUGUCUUCACUCCC | hsa-miR-149 | YP00204321 | {Ferracin, 2011 #133} |
| hsa-miR-152-3p | B08 | UCAGUGCAUGACAGAACUUGG | hsa-miR-152 | YP00204294 | {Rosenfeld, 2008 #21} |
| hsa-miR-181a-5p | B09 | AACAUUCAACGCUGUCGGUGAGU | hsa-miR-181a | YP00206081 | {Rosenfeld, 2008 #21} |
| hsa-miR-181a-2-3p | B10 | ACCACUGACCGUUGACUGUACC | hsa-miR-181a-2* | YP00204142 | {Ferracin, 2011 #133} |
| hsa-miR-181b-5p | B11 | AACAUUCAUUGCUGUCGGUGGGU | hsa-miR-181b | YP00204530 | {Rosenfeld, 2008 #21} |
| UniSp6 CP | B12 | | | YP00203954 | |
| hsa-miR-182-5p | C01 | UUUGGCAAUGGUAGAACUCACACU | hsa-miR-182 | YP00206070 | {Ferracin, 2011 #133} |
| hsa-miR-183-5p | C02 | UAUGGCACUGGUAGAAUUCACU | hsa-miR-183 | YP00206030 | {Ferracin, 2011 #133} |
| hsa-miR-187-3p | C03 | UCGUGUCUUGUGUUGCAGCCGG | hsa-miR-187 | YP00204018 | {Rosenfeld, 2008 #21} |
| hsa-miR-187-5p | C04 | GGCUACAACACAGGACCCGGGC | hsa-miR-187* | YP00205920 | {Ferracin, 2011 #133} |
| hsa-miR-192-5p | C05 | CUGACCUAUGAAUUGACAGCC | hsa-miR-192 | YP00204099 | {Ferracin, 2011 #133} |
| hsa-miR-193a-3p | C06 | AACUGGCCUACAAAGUCCCAGU | hsa-miR-193a-3p | YP00204591 | {Ferracin, 2011 #133} |
| hsa-miR-193b-3p | C07 | AACUGGCCCUCAAAGUCCCGCU | hsa-miR-193b | YP00204226 | {Rosenfeld, 2008 #21} |
| hsa-miR-194-5p | C08 | UGUAACAGCAACUCCAUGUGGA | hsa-miR-194 | YP00204080 | {Ferracin, 2011 #133} |
| hsa-miR-194-3p | C09 | CCAGUGGGGCUGCUGUUAUCUG | hsa-miR-194* | YP00204204 | {Ferracin, 2011 #133} |
| hsa-miR-196a-5p | C10 | UAGGUAGUUUCAUGUUGUUGGG | hsa-miR-196a | YP00204386 | {Rosenfeld, 2008 #21} |
| hsa-miR-19b-3p | C11 | UGUGCAAAUCCAUGCAAAACUGA | hsa-miR-19b | YP00204450 | {Rosenfeld, 2008 #21} |
| NTC | C12 | | | | |
| hsa-miR-200a-3p | D01 | UAACACUGUCUGGUAACGAUGU | hsa-miR-200a | YP00204707 | {Ferracin, 2011 #133} |
| hsa-miR-200a-5p | D02 | CAUCUUACCGGACAGUGCUGGA | hsa-miR-200a* | YP00206063 | {Ferracin, 2011 #133} |
| hsa-miR-200b-3p | D03 | UAAUACUGCCUGGUAAUGAUGA | hsa-miR-200b | YP00206071 | {Ferracin, 2011 #133} |
| hsa-miR-200b-5p | D04 | CAUCUUACUGGGCAGCAUUGGA | hsa-miR-200b* | YP00204144 | {Ferracin, 2011 #133} |
| hsa-miR-200c-3p | D05 | UAAUACUGCCGGGUAAUGAUGGA | hsa-miR-200c | YP00204482 | {Ferracin, 2011 #133} |
| hsa-miR-204-5p | D06 | UUCCCUUUGUCAUCCUAUGCCU | hsa-miR-204 | YP00206072 | {Ferracin, 2011 #133} |
| hsa-miR-205-5p | D07 | UCCUUCAUUCCACCGGAGUCUG | hsa-miR-205 | YP00204487 | {Ferracin, 2011 #133} |
| hsa-miR-210-3p | D08 | CUGUGCGUGUGACAGCGGCUGA | hsa-miR-210 | YP00204333 | {Ferracin, 2011 #133} |
| hsa-miR-211-5p | D09 | UUCCCUUUGUCAUCCUUCGCCU | hsa-miR-211 | YP00204009 | {Ferracin, 2011 #133} |
| hsa-miR-214-3p | D10 | ACAGCAGGCACAGACAGGCAGU | hsa-miR-214 | YP00204510 | {Rosenfeld, 2008 #21} |
| hsa-miR-215-5p | D11 | AUGACCUAUGAAUUGACAGAC | hsa-miR-215 | YP00204598 | {Ferracin, 2011 #133} |
| UniSp3 IPC | D12 | | | YP02119288 | |
| hsa-miR-27b-3p | E01 | UUCACAGUGGCUAAGUUCUGC | hsa-miR-27b | YP00205915 | {Rosenfeld, 2008 #21} |
| hsa-miR-29a-3p | E02 | UAGCACCAUCUGAAAUCGGUUA | hsa-miR-29a | YP00204698 | {Rosenfeld, 2008 #21} |
| hsa-miR-29b-3p | E03 | UAGCACCAUUUGAAAUCAGUGUU | hsa-miR-29b | YP00204679 | {Rosenfeld, 2008 #21} |
| hsa-miR-29c-3p | E04 | UAGCACCAUUUGAAAUCGGUUA | hsa-miR-29c | YP00204729 | {Rosenfeld, 2008 #21} |
| hsa-miR-30a-5p | E05 | UGUAAACAUCCUCGACUGGAAG | hsa-miR-30a* | YP00205695 | {Ferracin, 2011 #133} |
| hsa-miR-30a-3p | E06 | CUUUCAGUCGGAUGUUUGCAGC | hsa-miR-30a | YP00204457 | {Ferracin, 2011 #133} |
| hsa-miR-30c-5p | E07 | UGUAAACAUCCUACACUCUCAGC | hsa-miR-30c | YP00204783 | {Ferracin, 2011 #133} |
| hsa-miR-31-5p | E08 | AGGCAAGAUGCUGGCAUAGCU | hsa-miR-31 | YP00204236 | {Ferracin, 2011 #133} |
| hsa-miR-31-3p | E09 | UGCUAUGCCAACAUAUUGCCAU | hsa-miR-31* | YP00204079 | {Ferracin, 2011 #133} |
| hsa-miR-340-3p | E10 | UCCGUCUCAGUUACUUUAUAGC | hsa-miR-340* | YP00204250 | {Ferracin, 2011 #133} |
| hsa-miR-342-3p | E11 | UCUCACACAGAAAUCGCACCCGU | hsa-miR-342-3p | YP00205625 | {Ferracin, 2011 #133} |
| hsa-miR-345-5p | E12 | GCUGACUCCUAGUCCAGGGCUC | hsa-miR-345 | YP00206006 | {Rosenfeld, 2008 #21} |
| hsa-miR-34a-5p | F01 | UGGCAGUGUCUUAGCUGGUUGU | hsa-miR-34a | YP00204486 | {Rosenfeld, 2008 #21} |
| hsa-miR-34b-3p | F02 | CAAUCACUAACUCCACUGCCAU | hsa-miR-34b | YP00204005 | {Rosenfeld, 2008 #21} |
| hsa-miR-361-5p | F03 | UUAUCAGAAUCUCCAGGGGUAC | hsa-miR-361-5p | YP00206054 | {Ferracin, 2011 #133} |
| hsa-miR-363-3p | F04 | AAUUGCACGGUAUCCAUCUGUA | hsa-miR-363 | YP00204726 | {Ferracin, 2011 #133} |
| hsa-miR-372-3p | F05 | AAAGUGCUGCGACAUUUGAGCGU | hsa-miR-372 | YP00204137 | {Rosenfeld, 2008 #21} |
| hsa-miR-373-3p | F06 | GAAGUGCUUCGAUUUUGGGGUGU | hsa-miR-373 | YP00204604 | {Rosenfeld, 2008 #21} |
| hsa-miR-375-3p | F07 | UUUGUUCGUUCGGCUCGCGUGA | hsa-miR-375 | YP00204362 | {Ferracin, 2011 #133} |
| hsa-miR-382-5p | F08 | GAAGUUGUUCGUGGUGGAUUCG | hsa-miR-382 | YP00204169 | {Rosenfeld, 2008 #21} |
| hsa-miR-485-5p | F09 | AGAGGCUGGCCGUGAUGAAUUC | hsa-miR-485-5p | YP02112548 | {Ferracin, 2011 #133} |
| hsa-miR-506-3p | F10 | UAAGGCACCCUUCUGAGUAGA | hsa-miR-506 | YP00204539 | {Ferracin, 2011 #133} |
| hsa-miR-508-3p | F11 | UGAUUGUAGCCUUUUGGAGUAGA | hsa-miR-508-3p | YP00204480 | {Ferracin, 2011 #133} |
| hsa-miR-509-3p | F12 | UGAUUGGUACGUCUGUGGGUAG | hsa-miR-509-3p | YP00204458 | {Ferracin, 2011 #133} |
| hsa-miR-510-5p | G01 | UACUCAGGAGAGUGGCAAUCAC | hsa-miR-510 | YP00204349 | {Ferracin, 2011 #133} |
| hsa-miR-514a-3p | G02 | AUUGACACUUCUGUGAGUAGA | hsa-miR-514 | YP00205931 | {Ferracin, 2011 #133} |
| hsa-miR-552-3p | G03 | AACAGGUGACUGGUUAGACAA | hsa-miR-552 | YP00206032 | {Ferracin, 2011 #133} |
| hsa-miR-649 | G04 | AAACCUGUGUUGUUCAAGAGUC | hsa-miR-649 | YP00204146 | |
| hsa-miR-650 | G05 | AGGAGGCAGCGCUCUCAGGAC | hsa-miR-650 | YP00204233 | {Ferracin, 2011 #133} |

| | | | | | |
|---|---|---|---|---|---|
| hsa-miR-661 | G06 | UGCCUGGGUCUCUGGCCUGCGCGU | hsa-miR-661 | YP00204657 | {Rosenfeld, 2008 #21} |
| hsa-miR-873-5p | G07 | GCAGGAACUUGUGAGUCUCCU | hsa-miR-873 | YP00204175 | {Ferracin, 2011 #133} |
| hsa-miR-9-3p | G08 | AUAAAGCUAGAUAACCGAAAGU | hsa-miR-9* | YP00204620 | {Rosenfeld, 2008 #21} |
| hsa-miR-92a-3p | G09 | UAUUGCACUUGUCCCGGCCUGU | hsa-miR-92 | YP00204258 | {Rosenfeld, 2008 #21} |
| hsa-miR-92b-3p | G10 | UAUUGCACUCGUCCCGGCCUCC | hsa-miR-92b | YP00204384 | {Rosenfeld, 2008 #21} |
| hsa-miR-96-5p | G11 | UUUGGCACUAGCACAUUUUUGCU | hsa-miR-96 | YP00204417 | {Ferracin, 2011 #133} |
| hsa-miR-99a-5p | G12 | AACCCGUAGAUCCGAUCUUGUG | hsa-miR-99a | YP00204521 | {Rosenfeld, 2008 #21} |
| SNORD44 (hsa) | H01 | | | YP00203902 | |
| SNORD48 (hsa) | H02 | | | YP00203903 | |
| U6 snRNA (hsa) | H03 | | | YP00203907 | |
| hsa-miR-24-3p | H04 | UGGCUCAGUUCAGCAGGAACAG | hsa-miR-24 | YP00204260 | {Ferracin, 2015 #98;Ferracin, 2010 #41} |
| hsa-miR-21-5p | H05 | UAGCUUAUCAGACUGAUGUUGA | hsa-miR-21 | YP00204230 | {Rosenfeld, 2008 #21} |
| hsa-miR-16-5p | H06 | UAGCAGCACGUAAAUAUUGGCG | hsa-miR-16 | YP00205702 | {Ferracin, 2015 #98;Ferracin, 2010 #41} |
| hsa-miR-320a | H07 | AAAAGCUGGGUUGAGAGGGCGA | hsa-miR-320a | YP00206042 | {Ferracin, 2015 #98;Ferracin, 2010 #41} |
| hsa-miR-224-5p | H08 | UCAAGUCACUAGUGGUUCCGUUUAG | hsa-miR-224-5p | YP00204641 | {Ferracin, 2015 #98;Ferracin, 2010 #41} |
| hsa-miR-423-5p | H09 | UGAGGGGCAGAGAGCGAGACUUU | hsa-miR-423-5p | YP00205624 | {Ferracin, 2015 #98;Ferracin, 2010 #41} |
| hsa-miR-25-3p | H10 | CAUUGCACUUGUCUCGGUCUGA | hsa-miR-25 | YP00204361 | {Ferracin, 2015 #98;Ferracin, 2010 #41} |
| hsa-miR-331-3p | H11 | GCCCCUGGGCCUAUCCUAGAA | hsa-miR-331-3p | YP00206046 | {Ferracin, 2015 #98;Ferracin, 2010 #41} |
| hsa-miR-103a-3p | H12 | AGCAGCAUUGUACAGGGCUAUGA | hsa-miR-103-20 | YP00204063 | {Ferracin, 2015 #98;Ferracin, 2010 #41} |
