## Supplementary Table 3 for "Molecular diagnosis and prognosis of cancers of unknown-primary (CUPs): progress from a microRNA-based droplet digital PCR assay"

**Supplementary Table 3. Error rates of the PAMR and LASSO models for each tumor class.**

| Tumor class | PAMR | LASSO |
|---|---|---|
| BLCA | 0.75 | 0.75 |
| PAAD | 0.69 | 0.75 |
| TNBC | 0.75 | 0.67 |
| LUSC | 0.22 | 0.52 |
| LIHC | 0.37 | 0.48 |
| HNSC | 0.49 | 0.47 |
| OV | 0.75 | 0.45 |
| BRCA | 0.00 | 0.39 |
| CHOL | 0.50 | 0.38 |
| KICA | 0.25 | 0.29 |
| GI-NET | 0.20 | 0.28 |
| TGSC | 0.25 | 0.25 |
| STAD-CRC | 0.16 | 0.15 |
| SKCM | 0.16 | 0.06 |
| LUAD | 0.08 | 0.04 |
| UCEC | 0.18 | 0.03 |
| PRAD | 0.02 | 0.00 |
