## Supplementary Table 4 for "Molecular diagnosis and prognosis of cancers of unknown-primary (CUPs): progress from a microRNA-based droplet digital PCR assay"

**Supplementary Table 4. Primary site prediction in metastases of known origin.**

| Sample ID | Primary site | PAMR 1st prediction | PAMR 2nd prediction | LASSO 1st prediction | LASSO 2nd prediction |
|---|---|---|---|---|---|
| CB048 | BRCA | BRCA | PAAD | BRCA | STAD-CRC |
| CB044 | CRC | STAD-CRC | PAAD | STAD-CRC | LIHC |
| PF088 | UCEC | UCEC | PAAD | PAAD | PRAD |
| PF086 | STAD | STAD-CRC | PAAD | STAD-CRC | PAAD |
| CB073 | HNSC | TNBC | HNSC | TNBC | HNSC |
| PF095 | KICA | CHOL | LIHC | OV | PAAD |
| PF010 | LUAD | BRCA | LUAD | LUAD | PAAD |
| CB049 | PAAD | STAD-CRC | PAAD | STAD-CRC | PAAD |
| PF087 | PRAD | STAD-CRC | PAAD | STAD-CRC | PAAD |
| PF079 | SKCM | LUAD | SKCM | TNBC | KICA |
