## Supplementary Table 5 for "Molecular diagnosis and prognosis of cancers of unknown-primary (CUPs): progress from a microRNA-based droplet digital PCR assay"

**Supplementary Table 5 - CUP probabilities with PAMR and LASSO classifier models**

| ID sample | first_pamr | first_pamr_prob | second_pamr | second_pamr_prob | first_lasso | first_lasso_prob | second_lasso | second_lasso_prob |
|---|---|---|---|---|---|---|---|---|
| **CB002** | **BRCA** | **0.749** | UCEC | 0.105 | **STAD-CRC** | **0.276** | **BRCA** | **0.272** |
| **CB003** | **HNSC** | **0.506** | **LUSC** | **0.225** | **HNSC** | **0.288** | **LUSC** | **0.129** |
| **CB011** | GI-NET | 0.728 | LUAD | 0.157 | **PAAD** | **0.427** | GI-NET | 0.410 |
| **CB012** | **CHOL** | **0.987** | KICA | 0.010 | **LIHC** | **0.323** | **CHOL** | **0.224** |
| **CB013** | **STAD-CRC** | **0.921** | **PAAD** | **0.072** | **STAD-CRC** | **0.684** | **PAAD** | **0.113** |
| **CB014** | **TNBC** | **0.849** | LUAD | 0.128 | **TNBC** | **0.336** | CHOL | 0.260 |
| **CB033** | **BRCA** | **0.997** | STAD-CRC | 0.001 | **BRCA** | **0.338** | STAD-CRC | 0.275 |
| **CB053** | STAD-CRC | 0.719 | **BRCA** | **0.277** | STAD-CRC | 0.688 | **BRCA** | **0.103** |
| **CB054** | **BRCA** | **0.979** | BLCA | 0.008 | CHOL | 0.470 | **BRCA** | **0.153** |
| **CB055** | **CHOL** | **0.999** | TNBC | 0.001 | TNBC | 0.283 | LIHC | 0.208 |
| **CB061** | **BRCA** | **0.996** | PAAD | 0.001 | **BRCA** | **0.528** | KICA | 0.168 |
| **CB062** | BRCA | 0.324 | KICA | 0.165 | **CHOL** | **0.186** | KICA | 0.142 |
| **CB064** | TNBC | 0.440 | LUAD | 0.364 | **HNSC** | **0.223** | TNBC | 0.204 |
| **CB071** | STAD-CRC | 0.289 | **CHOL** | **0.280** | **CHOL** | **0.558** | KICA | 0.095 |
| **CB090** | **STAD-CRC** | **1.000** | PAAD | 0.000 | **STAD-CRC** | **0.871** | OV | 0.038 |
| **CB095** | **BLCA** | **0.601** | TNBC | 0.368 | **BLCA** | **0.513** | TNBC | 0.360 |
| **CB097** | **PAAD** | **0.782** | STAD-CRC | 0.139 | **PAAD** | **0.429** | STAD-CRC | 0.351 |
| **CB103** | **LUAD** | **0.963** | TNBC | 0.036 | **LUAD** | **0.653** | TNBC | 0.122 |
| **CB104** | **LUSC** | **0.996** | BLCA | 0.002 | **LUSC** | **0.414** | BLCA | 0.408 |
| **CB110** | **OV** | **0.901** | TNBC | 0.095 | **OV** | **0.865** | TNBC | 0.071 |
| **CB112** | **LUAD** | **0.790** | KICA | 0.025 | **LUAD** | **0.479** | KICA | 0.109 |
| **CB115** | **PAAD** | **0.455** | **STAD-CRC** | **0.253** | **STAD-CRC** | **0.174** | **PAAD** | **0.147** |
| **CB116** | **STAD-CRC** | **0.979** | **PAAD** | **0.020** | **STAD-CRC** | **0.767** | **PAAD** | **0.114** |
| **CB117** | **LUAD** | **0.878** | PAAD | 0.039 | **LUAD** | **0.272** | PAAD | 0.206 |
| **CB118** | **LUAD** | **0.994** | PAAD | 0.002 | **LUAD** | **0.598** | STAD-CRC | 0.128 |
| **CB125** | **STAD-CRC** | **0.961** | PAAD | 0.021 | **STAD-CRC** | **0.843** | LUAD | 0.031 |

| | | | | | | | | |
|---|---|---|---|---|---|---|---|---|
| **PF005** | **BRCA** | **0.958** | PAAD | 0.021 | **BRCA** | **0.186** | STAD-CRC | 0.183 |
| **PF006** | **LUAD** | **0.959** | BRCA | 0.018 | **LUAD** | **0.960** | PAAD | 0.012 |
| **PF007** | **STAD-CRC** | **0.982** | PAAD | 0.016 | **STAD-CRC** | **0.959** | TNBC | 0.012 |
| **PF011** | **LUSC** | **0.666** | HNSC | 0.165 | **LUSC** | **0.379** | PAAD | 0.111 |
| **PF013** | **CHOL** | **0.617** | KICA | 0.367 | **CHOL** | **0.103** | KICA | 0.092 |
| **PF017** | **PAAD** | **0.796** | **CHOL** | **0.180** | **PAAD** | **0.683** | **CHOL** | **0.052** |
| **PF018** | **STAD-CRC** | **0.435** | OV | 0.307 | **STAD-CRC** | **0.274** | OV | 0.249 |
| **PF019** | BRCA | 0.532 | **PAAD** | **0.383** | **PAAD** | **0.271** | BRCA | 0.165 |
| **PF020** | **PAAD** | **0.954** | STAD-CRC | 0.004 | **PAAD** | **0.577** | STAD-CRC | 0.118 |
| **PF021** | BRCA | 0.531 | PAAD | 0.361 | PAAD | 0.352 | STAD-CRC | 0.133 |
| **PF022** | **PAAD** | **0.977** | STAD-CRC | 0.014 | **PAAD** | **0.578** | STAD-CRC | 0.270 |
| **PF024** | **LUSC** | **0.661** | BRCA | 0.295 | **LUSC** | **0.219** | PAAD | 0.179 |
| **PF025** | **LIHC** | **1.000** | CHOL | 0.000 | **LIHC** | **0.975** | CHOL | 0.008 |
| **PF059** | **LUAD** | **0.884** | TNBC | 0.073 | STAD-CRC | 0.178 | **LUAD** | **0.164** |
| **PF080** | **PAAD** | **0.711** | LUAD | 0.259 | **PAAD** | **0.406** | **STAD-CRC** | **0.185** |
| **CB098** | **TNBC** | **0.584** | LUAD | 0.396 | **TNBC** | **0.435** | LUAD | 0.198 |
| **CB100** | LUAD | 0.727 | **TNBC** | **0.252** | **TNBC** | **0.386** | LUAD | 0.163 |
| **CB101** | **TNBC** | **0.973** | HNSC | 0.026 | **TNBC** | **0.722** | OV | 0.159 |
| **CB102** | **BRCA** | **0.849** | LUAD | 0.108 | **BRCA** | **0.203** | **TNBC** | **0.168** |
| **CB105** | **STAD-CRC** | **0.804** | PAAD | 0.162 | **STAD-CRC** | **0.506** | GI-NET | 0.204 |
| **CB106** | GI-NET | 0.842 | **STAD-CRC** | **0.098** | **STAD-CRC** | **0.567** | GI-NET | 0.203 |
| **CB108** | BRCA | 0.536 | LUAD | 0.235 | **STAD-CRC** | **0.210** | BRCA | 0.150 |
| **CB109** | **STAD-CRC** | **0.866** | PAAD | 0.107 | **STAD-CRC** | **0.700** | PAAD | 0.067 |
| **CB119** | **LUAD** | **0.416** | **HNSC** | **0.327** | **HNSC** | **0.305** | TNBC | 0.167 |
| **CB120** | **LUAD** | **0.665** | TNBC | 0.144 | **HNSC** | **0.270** | **LUAD** | **0.222** |
| **CB121** | **PAAD** | **0.394** | CHOL | 0.335 | LUAD | 0.349 | **CHOL** | **0.251** |
| **CB122** | **CHOL** | **0.741** | PAAD | 0.231 | **PAAD** | **0.316** | **CHOL** | **0.168** |
