## Supplementary Table 6 for "Molecular diagnosis and prognosis of cancers of unknown-primary (CUPs): progress from a microRNA-based droplet digital PCR assay"

**Supplementary table 6. Association of miRNA expression with CUP overall survival (all miRNAs)**

| miRNA | HR | lower 95% | upper 95% | *p*-value |
|---|---|---|---|---|
| miR-124-3p | 0.11 | 0.03 | 0.36 | **0.00** |
| miR-21-5p | 7.00 | 2.10 | 23.00 | **0.00** |
| miR-9-3p | 0.29 | 0.12 | 0.71 | **0.01** |
| miR-149-5p | 0.32 | 0.13 | 0.78 | **0.01** |
| miR-372-3p | 0.33 | 0.12 | 0.89 | **0.03** |
| miR-485-5p | 0.37 | 0.16 | 0.90 | **0.03** |
| miR-375 | 9.60 | 1.30 | 73.00 | **0.03** |
| miR-25-3p | 0.26 | 0.08 | 0.87 | **0.03** |
| miR-27b-3p | 2.60 | 1.10 | 6.10 | **0.03** |
| miR-181a-2-3p | 0.38 | 0.15 | 0.93 | **0.03** |
| miR-10b-5p | 0.35 | 0.13 | 0.93 | **0.04** |
| miR-96-5p | 2.50 | 1.00 | 6.20 | **0.04** |
| miR-423-5p | 3.50 | 1.00 | 12.00 | **0.04** |
| miR-214-3p | 2.60 | 1.00 | 6.60 | **0.05** |
| miR-196a-5p | 0.31 | 0.09 | 1.10 | 0.06 |
| miR-193a-3p | 0.15 | 0.02 | 1.10 | 0.07 |
| miR-345-5p | 0.36 | 0.12 | 1.10 | 0.07 |
| miR-211-5p | 0.43 | 0.17 | 1.10 | 0.08 |
| miR-509-3p | 0.48 | 0.22 | 1.10 | 0.08 |
| miR-224-5p | 2.20 | 0.89 | 5.50 | 0.09 |
| miR-510-5p | 0.28 | 0.07 | 1.20 | 0.09 |
| miR-649 | 0.45 | 0.18 | 1.10 | 0.09 |
| miR-210-3p | 2.20 | 0.87 | 5.60 | 0.10 |
| miR-361-5p | 0.50 | 0.22 | 1.10 | 0.10 |
| miR-152-3p | 2.00 | 0.84 | 4.70 | 0.12 |
| miR-30a-5p | 2.00 | 0.83 | 4.60 | 0.12 |
| miR-508-3p | 0.51 | 0.22 | 1.20 | 0.12 |
| miR-873-5p | 0.42 | 0.14 | 1.30 | 0.12 |
| miR-320a | 0.49 | 0.20 | 1.20 | 0.12 |
| miR-514a-3p | 0.39 | 0.12 | 1.30 | 0.13 |
| miR-29c-3p | 4.30 | 0.58 | 32.00 | 0.15 |
| miR-92b-3p | 0.53 | 0.23 | 1.30 | 0.15 |
| miR-99a-5p | 0.52 | 0.22 | 1.30 | 0.15 |
| miR-187-5p | 1.80 | 0.76 | 4.40 | 0.17 |
| miR-373-3p | 0.56 | 0.24 | 1.30 | 0.17 |
| miR-194-5p | 1.70 | 0.78 | 3.90 | 0.18 |
| miR-200c-3p | 2.70 | 0.63 | 12.00 | 0.18 |
| miR-182-5p | 0.58 | 0.26 | 1.30 | 0.19 |
| let-7i-5p | 2.60 | 0.60 | 11.00 | 0.21 |
| miR-200a-3p | 1.70 | 0.74 | 3.90 | 0.21 |
| miR-187-3p | 2.60 | 0.56 | 12.00 | 0.22 |
| miR-29a-3p | 1.60 | 0.74 | 3.70 | 0.23 |
| miR-31-5p | 1.80 | 0.69 | 4.60 | 0.23 |
| miR-148b-3p | 0.40 | 0.08 | 1.90 | 0.25 |

| | | | | |
|---|---|---|---|---|
| miR-34a-5p | 3.30 | 0.43 | 25.00 | 0.25 |
| miR-204-5p | 0.59 | 0.23 | 1.50 | 0.26 |
| miR-30a-3p | 1.90 | 0.63 | 5.60 | 0.26 |
| miR-183-5p | 0.63 | 0.28 | 1.40 | 0.27 |
| miR-205-5p | 0.64 | 0.28 | 1.40 | 0.28 |
| miR-331-3p | 0.61 | 0.24 | 1.50 | 0.28 |
| miR-650 | 1.60 | 0.67 | 3.90 | 0.29 |
| miR-194-3p | 0.52 | 0.15 | 1.80 | 0.30 |
| miR-135b-5p | 2.80 | 0.36 | 21.00 | 0.32 |
| miR-552-3p | 1.70 | 0.60 | 4.70 | 0.32 |
| let-7e-5p | 0.64 | 0.26 | 1.60 | 0.34 |
| miR-181b-5p | 0.66 | 0.28 | 1.60 | 0.34 |
| miR-661 | 0.65 | 0.27 | 1.60 | 0.34 |
| miR-103a-3p | 2.60 | 0.35 | 20.00 | 0.35 |
| miR-363-3p | 0.68 | 0.29 | 1.60 | 0.36 |
| miR-506-3p | 0.66 | 0.26 | 1.70 | 0.38 |
| miR-215-5p | 1.40 | 0.63 | 3.30 | 0.39 |
| miR-192-5p | 1.40 | 0.61 | 3.20 | 0.42 |
| miR-130a-3p | 1.40 | 0.59 | 3.30 | 0.44 |
| miR-141-3p | 1.40 | 0.59 | 3.20 | 0.45 |
| miR-342-3p | 0.72 | 0.31 | 1.70 | 0.45 |
| miR-10a-3p | 1.50 | 0.54 | 3.90 | 0.46 |
| miR-146a-5p | 1.40 | 0.57 | 3.40 | 0.47 |
| miR-16-5p | 0.75 | 0.33 | 1.70 | 0.49 |
| miR-30c-5p | 0.71 | 0.26 | 1.90 | 0.50 |
| miR-92a-3p | 0.76 | 0.33 | 1.80 | 0.52 |
| miR-193b-3p | 0.76 | 0.32 | 1.80 | 0.54 |
| miR-142-3p | 1.40 | 0.49 | 3.80 | 0.55 |
| miR-106b-5p | 0.64 | 0.14 | 2.90 | 0.56 |
| miR-200b-5p | 1.80 | 0.24 | 14.00 | 0.56 |
| miR-382-5p | 1.30 | 0.54 | 2.90 | 0.59 |
| miR-340-3p | 0.69 | 0.16 | 3.00 | 0.62 |
| miR-138-5p | 0.82 | 0.36 | 1.90 | 0.65 |
| miR-10a-5p | 1.20 | 0.52 | 2.70 | 0.69 |
| miR-34b-3p | 1.20 | 0.41 | 3.80 | 0.70 |
| miR-200b-3p | 1.20 | 0.51 | 2.60 | 0.71 |
| miR-29b-3p | 0.83 | 0.24 | 2.90 | 0.77 |
| miR-24-3p | 1.10 | 0.50 | 2.50 | 0.79 |
| miR-126-5p | 0.91 | 0.40 | 2.10 | 0.82 |
| miR-19b-3p | 0.94 | 0.41 | 2.20 | 0.89 |
| miR-31-3p | 1.00 | 0.46 | 2.40 | 0.93 |
| miR-145-5p | 1.00 | 0.40 | 2.70 | 0.94 |
| miR-181a-5p | 0.97 | 0.42 | 2.20 | 0.94 |
| miR-200a-5p | 0.98 | 0.44 | 2.20 | 0.96 |

For each miRNA the hazard ratio (HR) with 95% confidence interval and *p*-value (*p*) is reported for OS. Significant p-values (<0.05) are reported in bold.
